# A soluble ACE2 microbody protein fused to a single immunoglobulin Fc domain is a potent inhibitor of SARS-CoV-2 infection in cell culture

**DOI:** 10.1101/2020.09.16.300319

**Authors:** Takuya Tada, Chen Fan, Ramanjit Kaur, Kenneth A. Stapleford, Harry Gristick, Crina Nimigean, Nathaniel R. Landau

## Abstract

Soluble forms of ACE2 have recently been shown to inhibit SARS-CoV-2 infection. We report on an improved soluble ACE2, termed a “microbody” in which the ACE2 ectodomain is fused to Fc domain 3 of the immunoglobulin heavy chain. The protein is smaller than previously described ACE2-Ig Fc fusion proteins and contains an H345A mutation in the ACE2 catalytic active site that inactivates the enzyme without reducing its affinity for the SARS-CoV-2 spike. The disulfide-bonded ACE2 microbody protein inhibited entry of lentiviral SARS-CoV-2 spike protein pseudotyped virus and live SARS-CoV-2 with a potency 10-fold higher than unmodified soluble ACE2 and was active after initial virus binding to the cell. The ACE2 microbody inhibited the entry of ACE2-specific β coronaviruses and viruses with the high infectivity variant D614G spike. The ACE2 microbody may be a valuable therapeutic for COVID-19 that is active against SARS-CoV-2 variants and future coronaviruses that may arise.

## Introduction

As the severe acute respiratory syndrome coronavirus 2 (SARS-CoV-2) continues to spread worldwide, there is an urgent need for preventative vaccine and improved therapeutics for treatment of COVID-19. The development of therapeutic agents that block specific steps of the coronavirus replication cycle will be highly valuable both for treatment and prophylaxis. Coronavirus replication consists of attachment, uncoating, replication, translation, assembly and release, all of which are potential drug targets. Virus entry is particularly advantageous because as the first step in virus replication, it spares target cells from becoming infected and because drugs that block entry do not need to be cell permeable as the targets are externally exposed. In SARS-CoV-2 entry, the virus attaches to the target cell through the interaction of the spike glycoprotein (S) with its receptor, the angiotensin-converting enzyme 2 (ACE2) (Li, 2015; Li et al., 2005; Li et al., 2003), a plasma membrane protein carboxypeptidase that degrades angiotensin II to angiotensin-(1–7) [Ang-(1–7)] a vasodilator that promotes sodium transport in the regulation of cardiac function and blood pressure (Kuba et al., 2010; Riordan, 2003; Tikellis and Thomas, 2012). ACE2 binding triggers S protein-mediated fusion of the viral envelope with the cell plasma membrane or intracellular endosomal membranes. The S protein is synthesized as a single polypeptide that is cleaved by the cellular protease furin into S1 and S2 subunits in the endoplasmic reticulum and then further processed by TMPRSS2 on target cells (Glowacka et al., 2011; Hoffmann et al., 2020; Matsuyama et al., 2010; Shulla et al., 2011). The S1 subunit contains the receptor binding domain (RBD) which binds to ACE2 while S2 mediates virus-cell fusion (Belouzard et al., 2012; Fehr and Perlman, 2015; Heald-Sargent and Gallagher, 2012; Li et al., 2006; Shang et al., 2020). Cells that express ACE2 are potential targets of the virus. These include cells in the lungs, arteries, heart, kidney, and intestines (Harmer et al., 2002; Ksiazek et al., 2003; Leung et al., 2003).

The use of soluble receptors to prevent virus entry by competitively binding to viral envelope glycoproteins was first explored for HIV-1 with soluble CD4. In early studies, a soluble form of CD4 deleted for the transmembrane and cytoplasmic domains was found to block virus entry *in vitro* (Daar et al., 1990; Haim et al., 2009; Orloff et al., 1993; Schenten et al., 1999; Sullivan et al., 1998). Fusion of the protein to an immunoglobulin Fc region, termed an “immunoadhesin”, increased the avidity for gp120 by dimerizing the protein and served to increase the half-life of the protein *in vivo*. An enhanced soluble CD4-Ig containing a peptide derived from the HIV-1 coreceptor CCR5 was found to potently block infection and to protect rhesus macaques from infection (Chiang et al., 2012). The soluble receptor approach to blocking virus entry has been recently applied to SARS-CoV-2 through the use of recombinant human soluble ACE2 protein (hrsACE2) (Kuba et al., 2005; Monteil et al., 2020; Wysocki et al., 2010) or hrsACE2-IgG which encodes soluble ACE2 and the Fc region of the human immunoglobulin G (IgG) (Case et al., 2020; Lei et al., 2020; Qian and Hu, 2020) which were shown to inhibit of SARS-CoV and SARS-CoV-2 entry in a mouse model. In phase 1 and phase 2 clinical trials (Haschke et al., 2013; Khan et al., 2017), the protein showed partial antiviral activity but short half-live. Addition of the Fc region increased the half-life of the protein *in vivo*. A potential concern with the addition of the Ig Fc region is the possibility of enhancement, similar to what occurs with antibody-dependent enhancement in which anti-spike protein antibody attaches to Fc receptors on immune cells, facilitating infection rather than preventing it (Eroshenko et al., 2020).

We report, here on a soluble human ACE2 “microbody” in which the ACE2 ectodomain is fused to domain 3 of immunoglobulin G heavy chain Fc region (IgG-CH3) (Maute et al., 2015). The IgG-CH3 Fc domain served to dimerize the protein, increasing its affinity for the SARS-CoV-2 S and decreasing the molecular mass of the protein. The ACE2 microbody did not bind to cell surface Fc receptor, reducing any possibility of infection enhancement. Mutation of the active site H345 to alanine in the ACE2 microbody protein, a mutation that has been shown to inactivate ACE2 catalytic activity (Guy et al., 2005), did not decrease its affinity for the S protein. The dimeric ACE2 microbody had about 10-fold higher antiviral activity than soluble ACE2, which was also a dimer, and high a higher affinity for virion binding. The ACE2 microbody blocked virus entry into ACE2.293T cells that over-expressed ACE2 as well as all of the cell-lines tested and was fully active against the D614G variant spike protein and a panel of β coronavirus spike proteins.

## Results

### SARS-CoV-2 Δ19 S protein was incorporated in pseudotyped virion and had higher infectivity

As a means to study SARS-CoV-2 entry, we developed an assay based on SARS-CoV-2 S protein pseudotyped lentiviral reporter viruses. The viruses package a lentiviral vector genome that encodes nanoluciferase and GFP separated by a P2A self-processing peptide, providing a convenient means to titer the virus, and the ability to use two different assays to measure infection. To pseudotype the virions, we constructed expression vectors for the full-length SARS-CoV-2 S and for a Δ19 variant deleted for the carboxy-terminal 19 amino acids that removes a reported endoplasmic reticulum retention sequence that blocks transit of the S protein to the cell surface (Giroglou et al., 2004) (**Figure 1A**). The vectors were constructed with or without a carboxy-terminal hemagglutinin (HA) epitope tag. Pseudotyped viruses were produced in 293T cells cotransfected with the dual nanoluciferase/GFP reporter lentiviral vector pLenti.GFP.NLuc, Gag/Pol expression vector pMDL and full-length S protein, the Δ19 S protein, vesicular stomatitis virus G protein (VSV-G) expression vector or without an envelope glycoprotein expression vector. Immunoblot analysis showed that full-length and Δ19 S proteins were expressed and processed into the cleaved S2 protein (S1 is not visible as it lacks an epitope tag). Analysis of the virions showed that the Δ19 S protein was packaged into virions at >20-fold higher levels than the full-length protein (**Figure 1B**). This difference was not the result of differences in virion production as similar amounts of virion p24 were present in the cell supernatant. Analysis of the transfected

**Figure 1.**
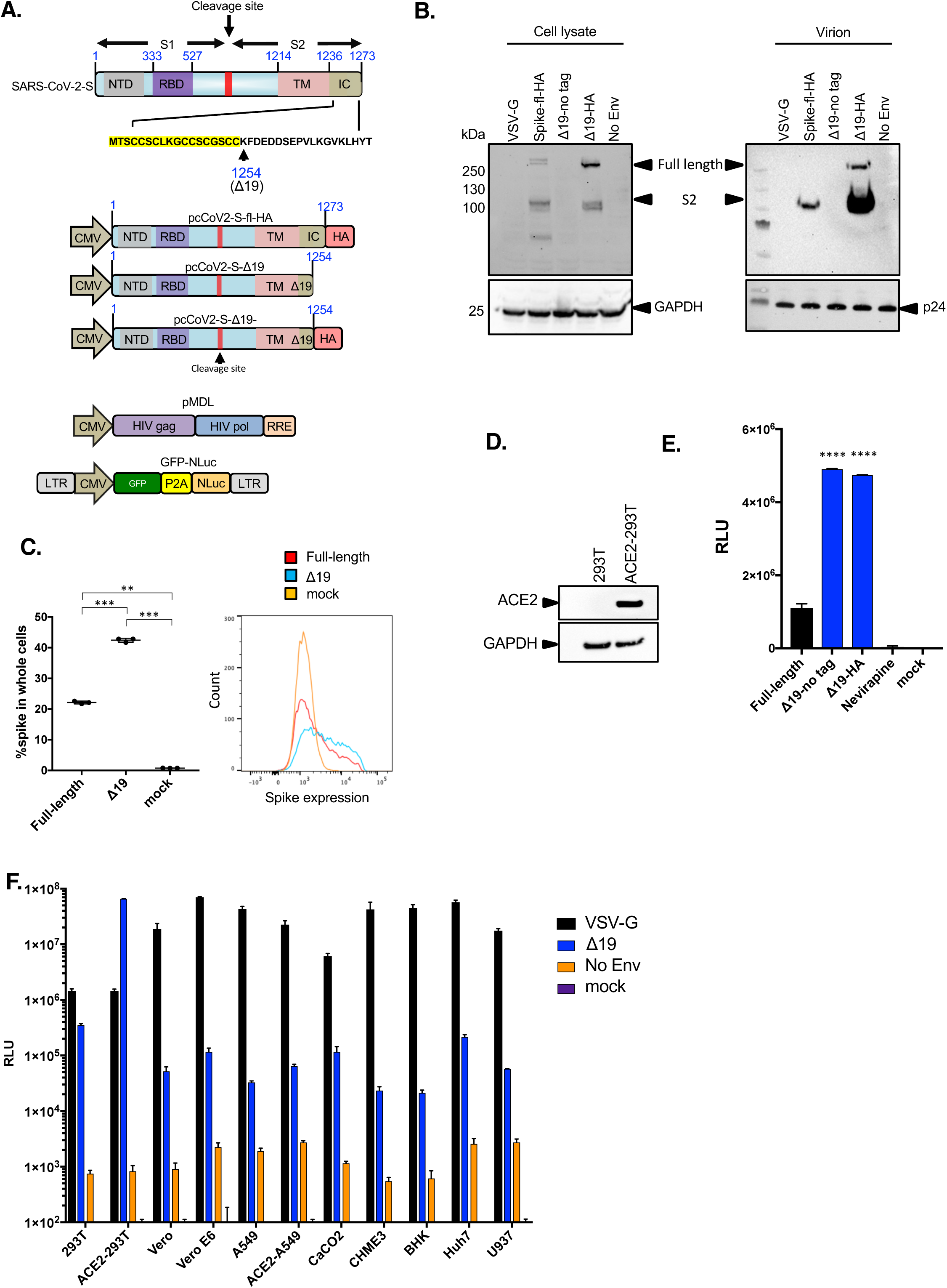
Δ19 SARS-CoV-2 pseudotyped lentiviral virion infection of ACE2.293T cells and ACE2 expressing cell-lines. (A) The domain structure of the SARS-CoV-2 S protein is diagrammed above. Yellow shading indicates the amino acids of the cytoplasmic domain retained following truncation of the 19 carboxy-terminal amino acids (Δ19). Vectors encoding full-length (fl) codon-optimized SARS-CoV-2 S protein or truncated Δ19 S protein, with or without a carboxy-terminal HA-tag and the dual nanoluciferase/GFP lentiviral vector pLenti.NLuc.GFP used to generate pseudotyped lentiviral particles are diagrammed (below). NTD: N-Terminal domain, RBD: Receptor-binding domain, TM: Transmembrane domain IC: intracellular domain, RRE: Rev response element, LTR: Long terminal repeat. (B) SARS-CoV-2 S proteins on pseudotyped lentiviral virions were analyzed. Transfected producer cell lysates (left) and supernatant virions (right) were analyzed on an immunoblot probed with anti-HA antibody. Cell lysate blots were probed with anti-GAPDH to normalize protein loading and virion blots were probed for HIV-1 p24 to normalize for virions. (C) 293T cells transfected with SARS-CoV-2 S protein expression vectors were analyzed by flow cytometry to detect the protein at the plasma membrane. (D) HA-tagged ACE2 expressed in control transfected 293T and clonal ACE2.293T cells were analyzed on an immunoblot probed with anti-HA antibody. (E) ACE2.293T cells were infected with virus pseudotyped by full-length or SARS-CoV-2 Δ19 S proteins. Two days post-infection, infectivity was measured by luciferase assay. The reverse transcriptase inhibitor nevirapine was added to one sample to control for free luciferase enzyme contamination of the virus stock. (F) A panel of cell-lines were infected with VSV-G, SARS-CoV-2 Δ19 S protein or no envelope (no Env) pseudotyped lentivirus. Luciferase activity was measured two days post-infection. The data are represented as the mean of triplicates ± the standard deviation. Statistical significance was calculated with the student-t test.

293T cells by flow cytometry showed a minor increase in the amount of cell surface Δ19 S protein as compared to full-length (**Figure 1C**) suggesting that deletion of the endoplasmic reticulum retention signal was not the primary cause of the increased virion packaging of the Δ19 S protein and may result from an inhibitory effect of the S protein cytoplasmic tail on virion incorporation. As a suitable target cell-line, we established a clonal, stably transfected 293T cell-line that expressed high levels of ACE2 (**Figures 1D and S1**). A comparison of the infectivity of the viruses on ACE2.293T cells showed that the Δ19 S protein pseudotype was about 2.5-fold more infectious than the full-length S protein pseudotype (**Figure 1E**). The HA-tag had no effect on infectivity and the nevirapine control demonstrated that the luciferase activity was the result of bona fide infection and not carried-over luciferase in the virus-containing supernatant.

To determine the cell-type tropism of the pseudotyped virus, we tested several standard laboratory cell-lines for susceptibility to infection to the Δ19 S protein pseudotyped virus. The VSV-G pseudotype, which has very high infectivity on most cell-types was tested for comparison and virus lacking a glycoprotein was included to control for potential receptor-independent virus uptake. The results showed high infectivity of the Δ19 S protein pseudotyped virus on ACE2.293T cells, intermediate infectivity on 293T, Vero, Vero E6, A549, ACE2.A549, CaCO2 and Huh7 and low infectivity on A549, CHME3, BHK and U937 (**Figure 1F**). Analysis by flow cytometry of cell surface ACE2 levels showed high level expression on ACE2.293T, intermediate levels expression on ACE2.A549 and low to undetectable levels on A549, CaCO2 and Huh7 (**Figure S1**). The low level of ACE2 expression on cells such as Vero and CaCO2 suggests that virus can use very small amounts of the receptor for entry. Moreover, the pseudotyped virus is a highly sensitive means with which to detect virus entry.

### An ACE2 microbody potently inhibits SARS-CoV-2 S-mediated virus entry

Soluble ACE2 and ACE2-Fc fusions have been shown to inhibit SARS-CoV-2 infection (Case et al., 2020; Kuba et al., 2010; Lei et al., 2020; Monteil et al., 2020; Qian and Hu, 2020; Wysocki et al., 2010). To increase the effectiveness of soluble ACE2 and improve therapeutic potential, we generated an ACE2-“microbody” in which the ACE2 ectodomain was fused to a single IgG CH3 domain of the IgG Fc region (**Figure 2A**). This domain contains the disulfide bonding cysteine residues of the IgG Fc that are required to dimerize the protein, which would serve to increase the ACE2 microbody avidity for ACE2. To prevent potential unwanted effects of the protein on blood pressure due to the catalytic activity of ACE2, we mutated H345, one of the key active site histidine residues of ACE2, to alanine, a mutation that has been shown to block catalytic activity (Guy et al., 2005). H345 lies underneath the S protein interaction site so was not predicted to interfere with S protein binding (**Figure 2B**). For comparison, we constructed vector encoding soluble ACE2 without the IgG CH3. The proteins were purified from transfected 293T cells and purified to homogeneity by Ni-NTA agarose affinity chromatography followed by size exclusion chromatography (**Figure S2**). The oligomerization state of the proteins was analyzed by SDS-PAGE under nonreducing and reducing conditions. Under reducing conditions, the ACE2 and ACE2.H345A microbody proteins and soluble ACE2 ran at the 130 kDa and 120 kDa, consistent with their calculated molecular mass (**Figures 2C** and S2). Under nonreducing conditions, the ACE2 microbody and ACE2.H345A microbody proteins ran at 250 kDa, consistent with dimers while the soluble ACE2 ran as a monomer with a mass of 120 kDa (**Figure 2C**). Analysis of the proteins by size-exclusion chromatography coupled with multi-angle light scattering (SEC-MALS) under nondenaturing conditions showed all three proteins to have a molecular mass consistent with dimers (**Figure 2D**). The mass of the ACE2 and ACE2.H345A microbody proteins was 218 kDa and 230 kDa, respectively, while soluble ACE2 was 180 kDa. Taken together, the results suggest that the ACE2 microbody proteins are disulfide-bonded dimers while soluble ACE2 is a nondisulfide-bonded dimer.

**Figure 2.**
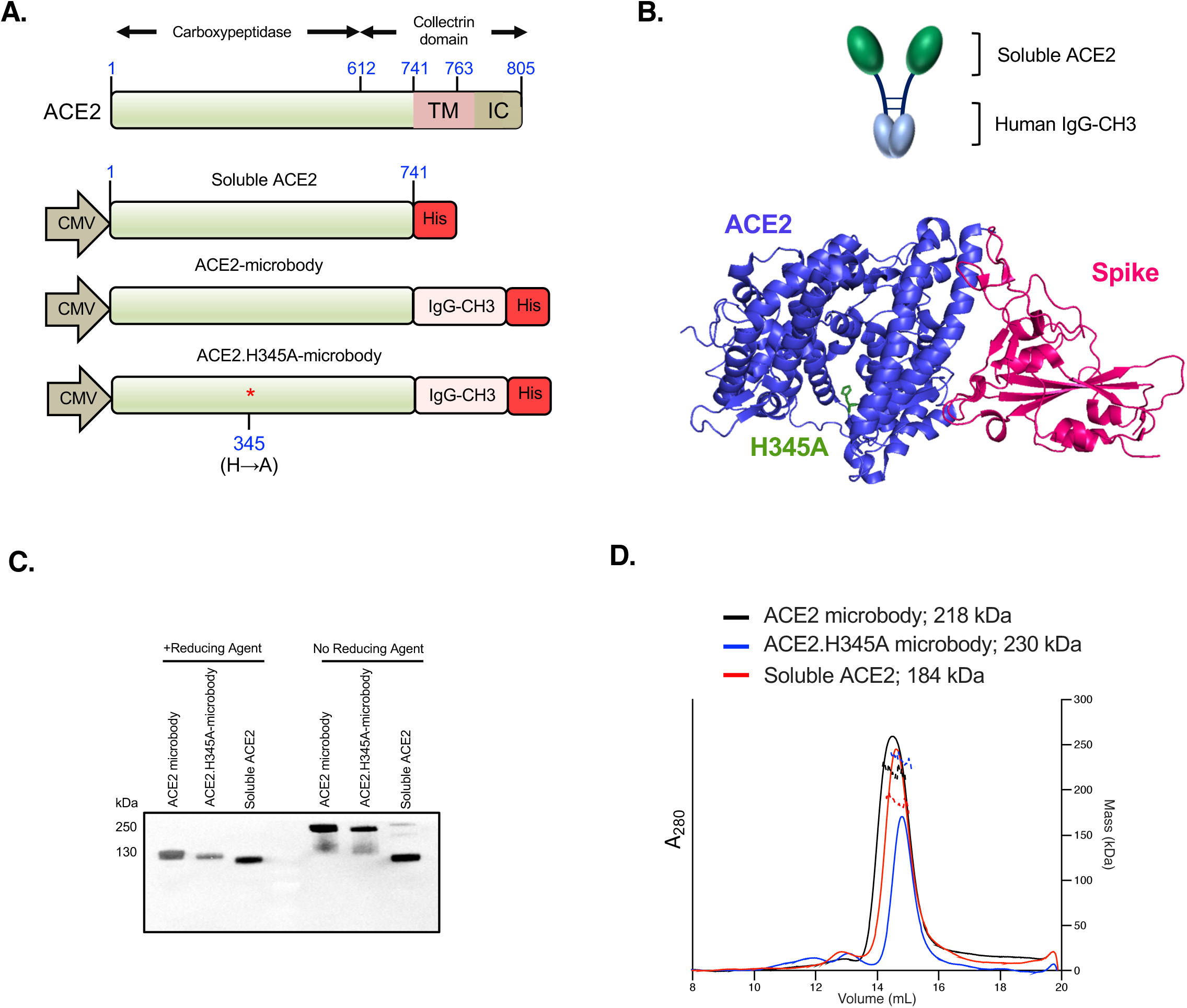
Wild-type and H345A ACE2 microbody proteins are disulfide bonded dimers. (A) The domains of ACE2 are shown with the structures of the soluble ACE2 (sACE2), ACE2 microbody and ACE2.H345A microbody proteins below. The soluble ACE2 proteins are deleted for the transmembrane (TM) and cytoplasmic domains. The ACE2 microbody proteins are fused to the human IgG CH3 domain each with a carboxy-terminal 8XHis-tag. IC: intracellular domain. (B) The diagram above shows the predicted dimeric structure of the ACE2 microbody protein. The 3D structure of the ACE2:spike protein complex was generated using Chimera software(Pettersen et al., 2004) using published coordinates(Lan et al., 2020). The position of the conserved active site H345 in the ACE2 carboxypeptidase domain is shown lying underneath the ACE2 interaction site. (C) 293T cells were transfected with sACE2, ACE2 microbody and ACE2.H345A microbody expression vectors. The proteins were pulled-down on NTA agarose beads and analyzed under reducing and nonreducing conditions on an immunoblot probed with anti-His-tag antibody. (D) The soluble ACE2 proteins were purified by metal chelate chromatography and size exclusion chromatography (SEC). The oligomerization state was determined by SEC multi-angle light scattering. The calculated molecular mass of each is shown.

### ACE2 microbody binds to SARS-CoV-2 pseudotyped virus

To compare the relative ability of the soluble ACE2 proteins to virions that display the S protein, we established a virion pull-down assay. Ni-NTA beads were incubated with a serial dilution of the carboxy-terminal His-tagged soluble ACE2 proteins. Free spike protein was removed and the beads were then incubated with a fixed amount of lentiviral pseudotyped virions. Free virions were removed and the bound virions were quantified by immunoblot analysis for virion p24 capsid protein. To confirm that virus binding to the beads was specific for the bead-bound ACE2, control virions lacking the spike were tested. The results showed that S protein pseudotyped virions bound to the beads while virions that lacked the S protein failed to bind, confirming that the binding was specific (**Figure 3A**). In addition, a high titer human serum from a recovered individual blocked binding of the virions to the bead-bound ACE2 microbody (**Figure S5**). Analysis of the soluble ACE2 and ACE2 microbody proteins that had bound to the beads showed that similar amounts of each proteins had bound (**Figure 3B**). Immunoblot analysis of virion binding to the bead-bound soluble ACE2 proteins showed that the wild-type and H345A microbody proteins both bound to virions more efficiently than soluble ACE2 (**Figure 3C**) and that the ACE2.H345A microbody bound more virions than the wild-type microbody protein. This was unexpected as H345 does not lie in the interaction surface with the S protein.

**Figure 3.**
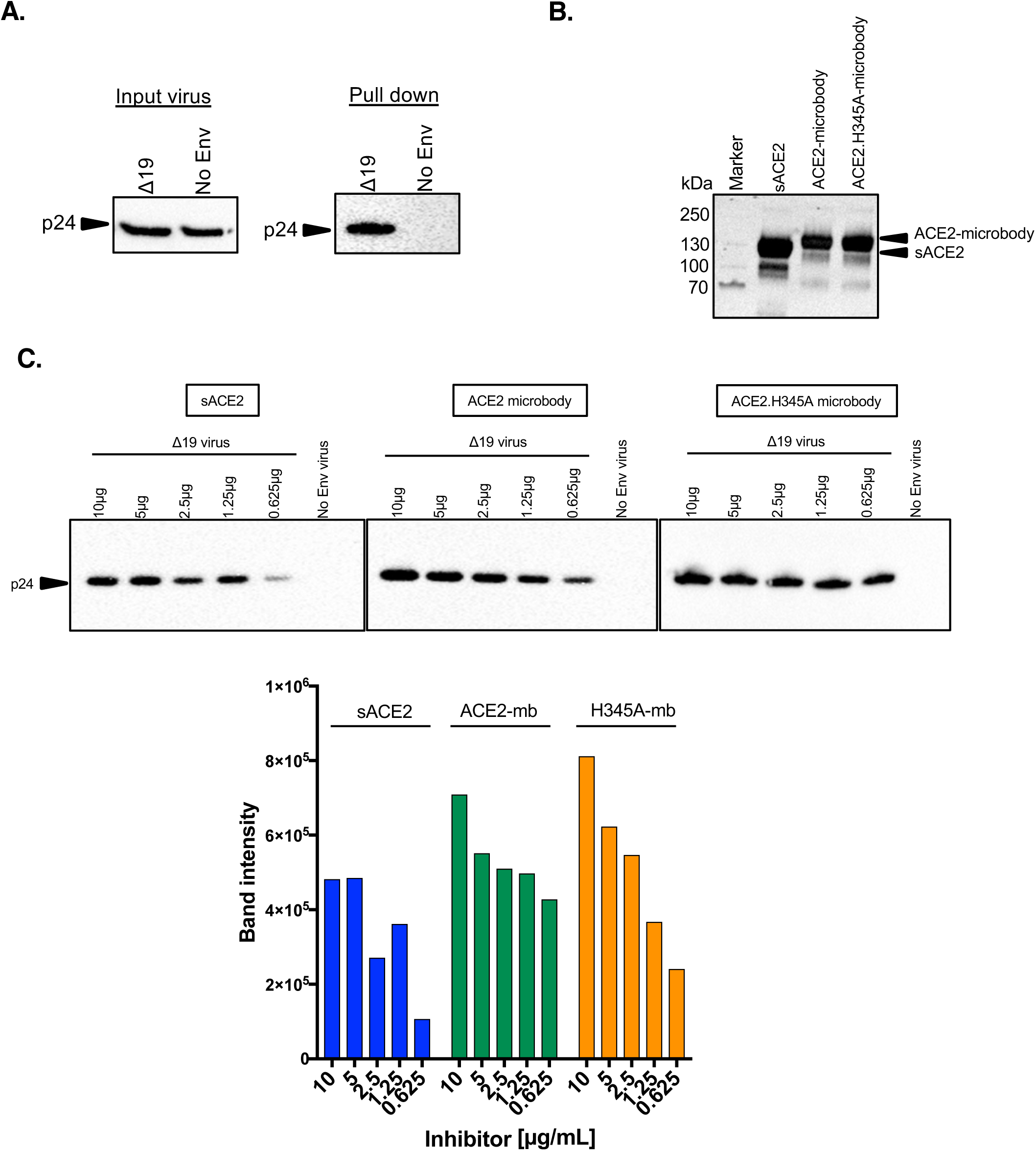
ACE2 microbody proteins bind with high affinity to SARS-CoV-2 S pseudotyped virions. Nickel agarose beads were coated for 1 hour with 10 μg of soluble ACE2 proteins. Unbound protein was removed and SARS-CoV-2 Δ19 S pseudotyped virions or virions lacking S protein were incubated with the beads. After 1 hour, unbound virions were removed and the bound virions were analyzed on an immunoblot probed with antibody p24 antibody. (A) Input virions and bead-bound virions were analyzed on an immunoblot probed with anti-p24 antibody. (B) Soluble ACE2 proteins bound to the nickel agarose beads were analyzed on an immunoblot probed with anti-His-tag antibody. (C) Soluble ACE2, wild-type ACE2 microbody and ACE2.H345A microbody proteins were serially diluted and bound to nickel agarose beads. The amount of bound virions was determined by immunoblot analysis with anti-p24 antibody. Quantification of the band intensities from the immunoblot is graphed below for soluble ACE2 (sACE2), wild-type ACE2 microbody (ACE2-mb) and ACE2.H345A microbody (H345A-mb).

### ACE2 microbody blocks SARS-CoV-2 pseudotyped virus infection

To determine the relative antiviral activity of soluble ACE2 and the ACE2 microbody proteins, we tested their ability to block the infection SARS-CoV-2 Δ19 S protein pseudotyped GFP/luciferase reporter virus. A fixed amount of pseudotyped reporter virus was incubated with the ACE2 proteins and then used to infect ACE2.293T cells. After 2 days, luciferase activity and the number of GFP+ cells in the infected cultures were analyzed. For comparison, a high titered recovered patient serum with a neutralizing titer of 1:330 (**figure S5**) was also tested. The results showed that soluble ACE2 had moderate inhibitory activity with an EC50 of 1.24 μg/ml. The ACE2 microbody was significantly more potent, with an EC50 of 0.36 μg/ml and the ACE2.H345A microbody protein was somewhat more potent than the wild-type ACE2 microbody with an EC50 of 0.15 μg/ml (**Figure 4A**). Inhibition of infection by the soluble ACE2 proteins was comparable to recovered patient serum although it is not possible to directly compare the two inhibitors as the mass amount of anti-S protein antibody in the serum is not known. To confirm the results, we analyzed the infected cells by flow cytometry to determine the number of GFP+ cells. The inhibition curves were similar to the luciferase curves, confirming that the ACE2 proteins had decreased the number of cells infected and did not simply reduce expression of the reporter protein (**Figure 4B, top**). Representative images of the GFP+ cells provide visual confirmation of the results (**Figure 4B, below)**. The inhibitory activity of the soluble ACE2 proteins was specific for the SARS-CoV-2 S protein as they did not inhibit VSV-G pseudotyped virus (**Figure 4C**). The ACE2 microbody was somewhat more active when tested on untransfected 293T that express low levels of ACE2 (**Figure 4D**).

**Figure 4.**
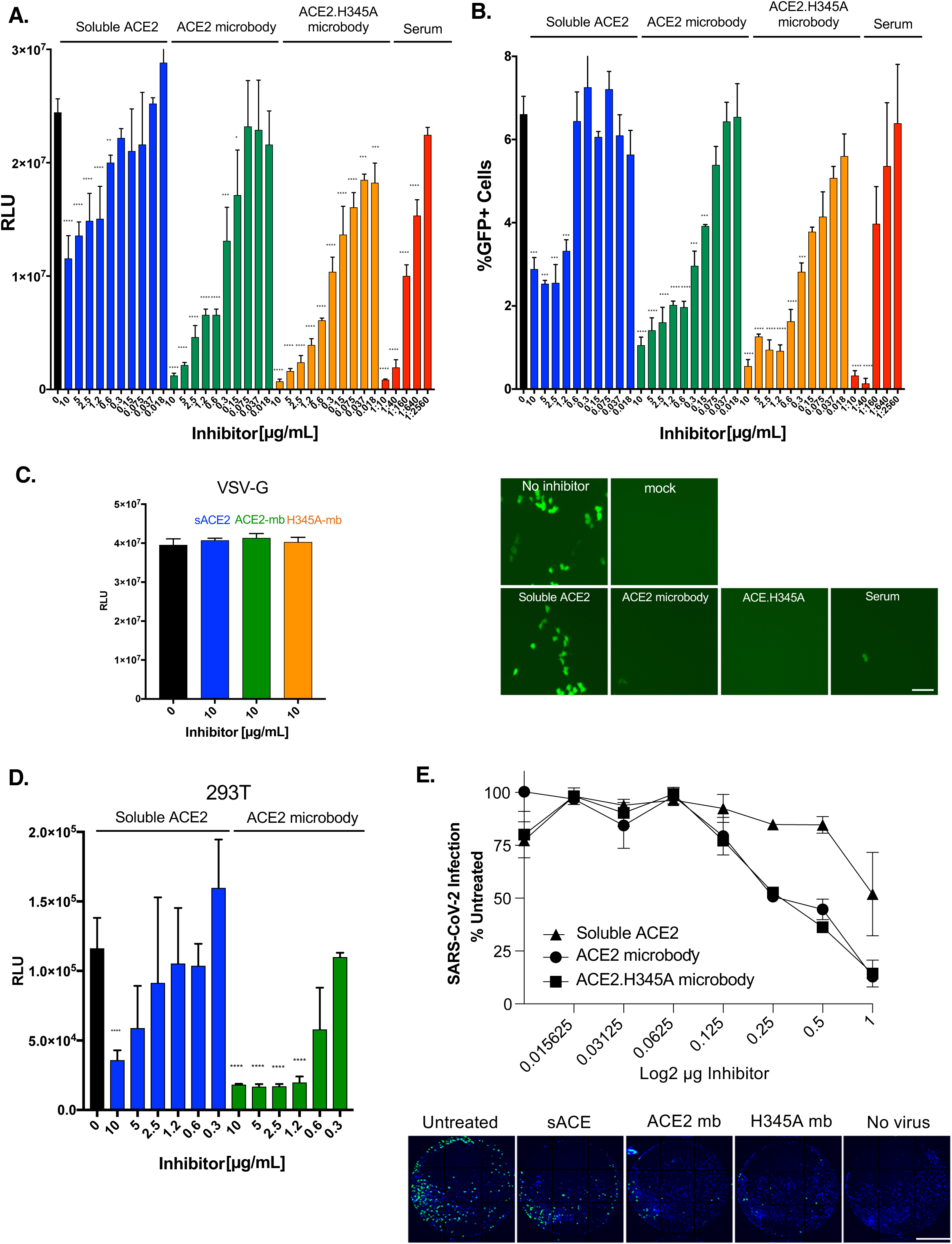
ACE2 and ACE2.H345A microbodies potently block virus entry and are active on different cell-lines. (A) Serially diluted soluble ACE2, ACE2 and ACE2.H345A microbody proteins were incubated for 30 min with SARS-CoV-2 Δ19.S-pseudotyped virus and then added to ACE2.293T cells. Luciferase activity was measured 2 days post-infection. For comparison, serially diluted convalescent COVID-19 patient serum was similarly analyzed. (B) The number of cells infected was determined by flow cytometry to quantify the GFP+ cells. The data are displayed as the percent GFP+ cells. Representative fluorescence microscopy images of the infected cells are shown below. Scale bar = 50 μm. (C) VSV-G pseudotyped lentiviral virions were incubated for 30 min with 10 μg/ml of soluble ACE2 proteins and then added to ACE2.293T cells. Luciferase activity was measured 2 days post-infection. (D) Δ19 S protein pseudotyped virus was incubated with serially diluted soluble ACE2 proteins for 30 min and then added 293T cells. (E) Serially diluted soluble ACE2 proteins were incubated with mNeonGreen SARS-CoV-2 for 30 min. and then added to ACE2.293T cells. After 24 hours, the GFP+ cells were counted. Fluorescent microscopy images of representative fields from wells treated with 1 μg soluble ACE2 and ACE2 microbody proteins are shown below. Scale bar = 2.1 mm. The data are displayed as the mean ± SD and significance is determined by student-t tests.

**Figure 5.**
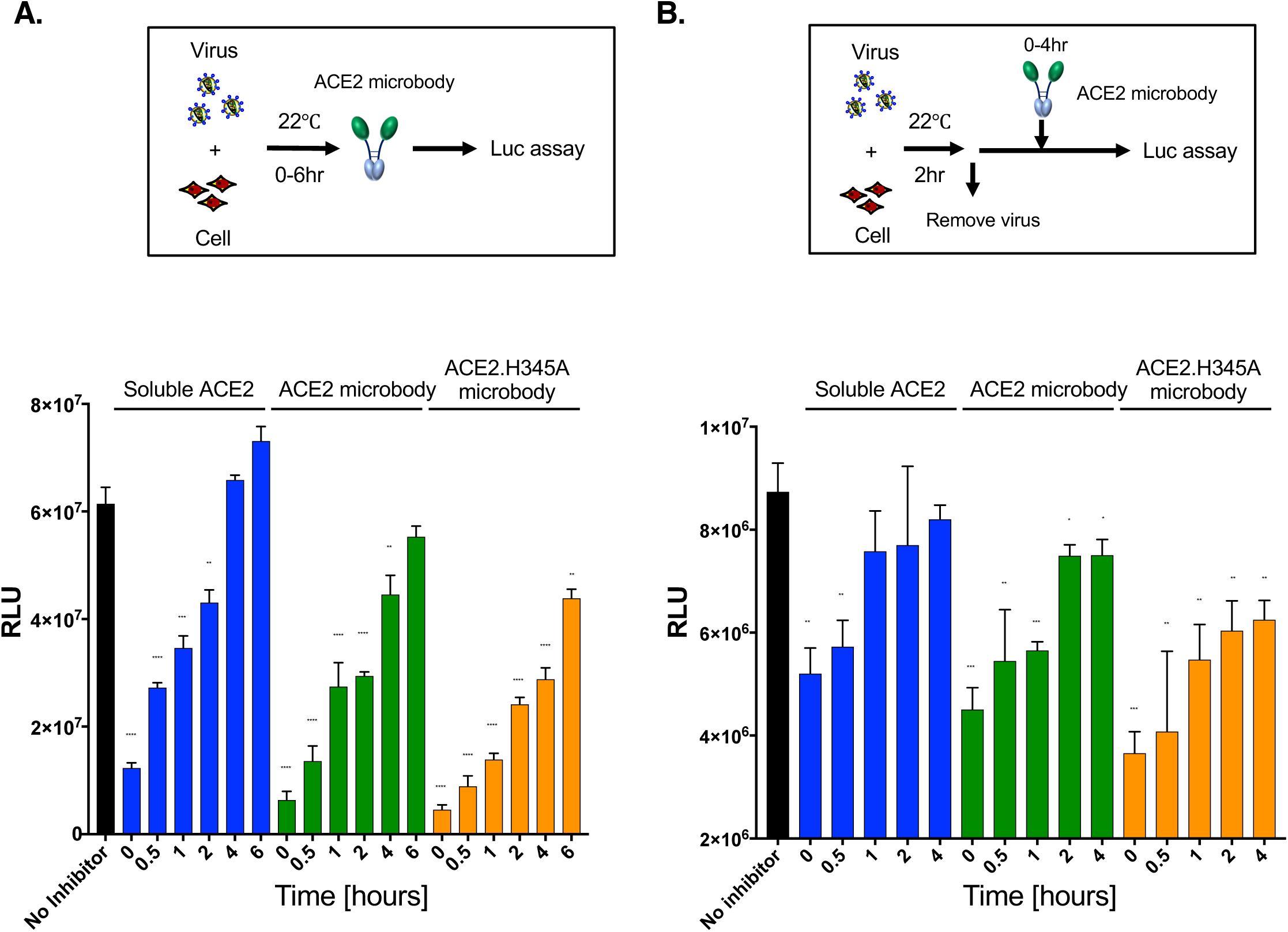
ACE2 microbody can act post-virus:cell binding. The kinetics of ACE2 microbody inhibition were analyzed in an escape from inhibition escape assay. (A) The experimental scheme is diagrammed above. SARS-CoV-2 Δ19.S-pseudotyped virus was added to ACE2.293T cells. Soluble ACE2 proteins were added immediately or at time points up to 6 hours later. Luciferase activity was measured 2 days post-infection. (B) As diagrammed above, virus was bound to target cells for 2 hours at 22°C and unbound virus was then removed. Soluble ACE2 proteins were added as in (A). The data are displayed as the mean ± SD and statistical significance determined by student-t tests.

To determine the ability of the ACE2 microbody proteins to block the replication of live SARS-CoV-2, we used the replication-competent SARS-CoV-2, icSARS-CoV-2mNG that encodes an mNeonGreen reporter gene in ORF7 (Xie et al., 2020). Serially diluted ACE2 microbody proteins were incubated with the virus and the mixture was then used to infect ACE2.293T cells. The results showed that 1-0.125 μg of ACE2 microbody protein blocked live virus replication (**Figure 4E**). Soluble ACE2 was less active; 1 μg of the protein had a 50% antiviral effect and the activity was lost with 0.5 μg. The antiviral activity of ACE2 proteins against live virus was similar to pseudotyped virus, except that in the live virus assay, the wild-type and H345A microbodies were of similar potency.

In the experiments described above, the proteins were incubated with virus prior to infection. To determine whether they would be active when at later time points, the ACE2 proteins were tested in an “escape from inhibition” assay in which the soluble ACE2 and ACE2 microbody proteins were added to cells at the same time as virus or up to 6 hours post-infection. The results showed that addition of the microbody together with the virus (t0) blocked the infection by 80%. Addition of the microbody 30 minutes post-infection maintained most of the antiviral effect, and even 2 hours post-infection the inhibitor blocked 55% of the infection. At 4 hours post-infection, the ACE2 microbody retained its blocking activity at 10 μg/ml but was less active with decreasing amounts of inhibitor (**Figure 5A**). These results suggest that the ACE2 microbody is highly efficient at neutralizing the virus when present before the virus has had a chance to bind to the cell and that it maintains its ability to block infection when added together with the cells and even 2 hours after the virus has been exposed to cells, a time which most of the virus has not yet bound to the cell.

To determine whether the ACE2 microbody could prevent virus entry once the virus bound to the cell, the virus was prebound by incubating it with cells for 1 hour at 22°C, the unbound virus was removed and the ACE2 microbody was added at increasing time points. The results showed that removal of the unbound virus after 1 hour incubation resulted in less infection as compared to when the virus was incubated with the cells for 4 hours, indicating that only a fraction of the virus had bound to cells. However, virus that was bound could be blocked by the ACE2 microbody for another 30 minutes post-binding (**Figure 5B**). The ability to block entry of the cell-bound virus suggests that virus binding results from a small number of spike molecules binding to ACE2. Over the next 30-60 minutes, additional spike:ACE2 interactions form, escaping the ability of the ACE2 microbody to block virus entry. The results demonstrate that the ACE2 microbody is a highly potent inhibitor of free virus and maintains its antiviral activity against virus newly-bound to the cell.

### ACE2 microbody blocks entry of virus with D614G mutant spike

A variant SARS-CoV-2 containing a D614G point mutation in the S protein has been found to be circulating in the human population with increasing prevalence (Daniloski et al., 2020; Eaaswarkhanth et al., 2020; Korber et al., 2020; Zhang et al., 2020). The D614G mutation was found to decrease shedding of the spike protein from the virus and to assume a fusion-ready conformation, resulting in increased infectivity and most likely contributing to its increasing prevalence. To determine the ability of the soluble ACE2 proteins to block entry of virus with the D614G S protein, we introduced the mutation into the Δ19 S protein expression vector and generated pseudotyped reporter viruses (**Figure 6A**). Analysis of the infectivity of the D614G and wild-type pseudotyped viruses on the panel of cell-lines showed that the mutation increased the infectivity of virus 2-4 fold on 293T, ACE2.293T, Vero and VeroE6 cells, consistent with previous reports (Daniloski et al., 2020; Yurkovetskiy et al., 2020; Zhang et al., 2020). Infectivity of the mutated virus was also increased in A549, ACE2.A549 CaCO2 although the overall infectivity of these cells was low (**Figure.6B)**. To determine the ability of the soluble ACE2 proteins to neutralize the virus with the variant S protein, serial dilutions of the soluble ACE2 proteins were tested for their ability to block wild-type and D614G S pseudotyped virus. The results showed that soluble ACE2 had moderate antiviral activity against wild-type virus, while the wild-type and H345A microbody proteins were more potent (**Figure 6C**). The ACE2.H345A microbody was somewhat more active at low concentrations than the wild-type protein. To test the relative binding affinity of the soluble ACE2 proteins for wild-type and D614G mutated spike, we tested the pseudotyped virions in the ACE2-virus binding assay (**Figure 6D**). The results showed that virus with the D614G S bound efficiently to soluble ACE2.

**Figure 6.**
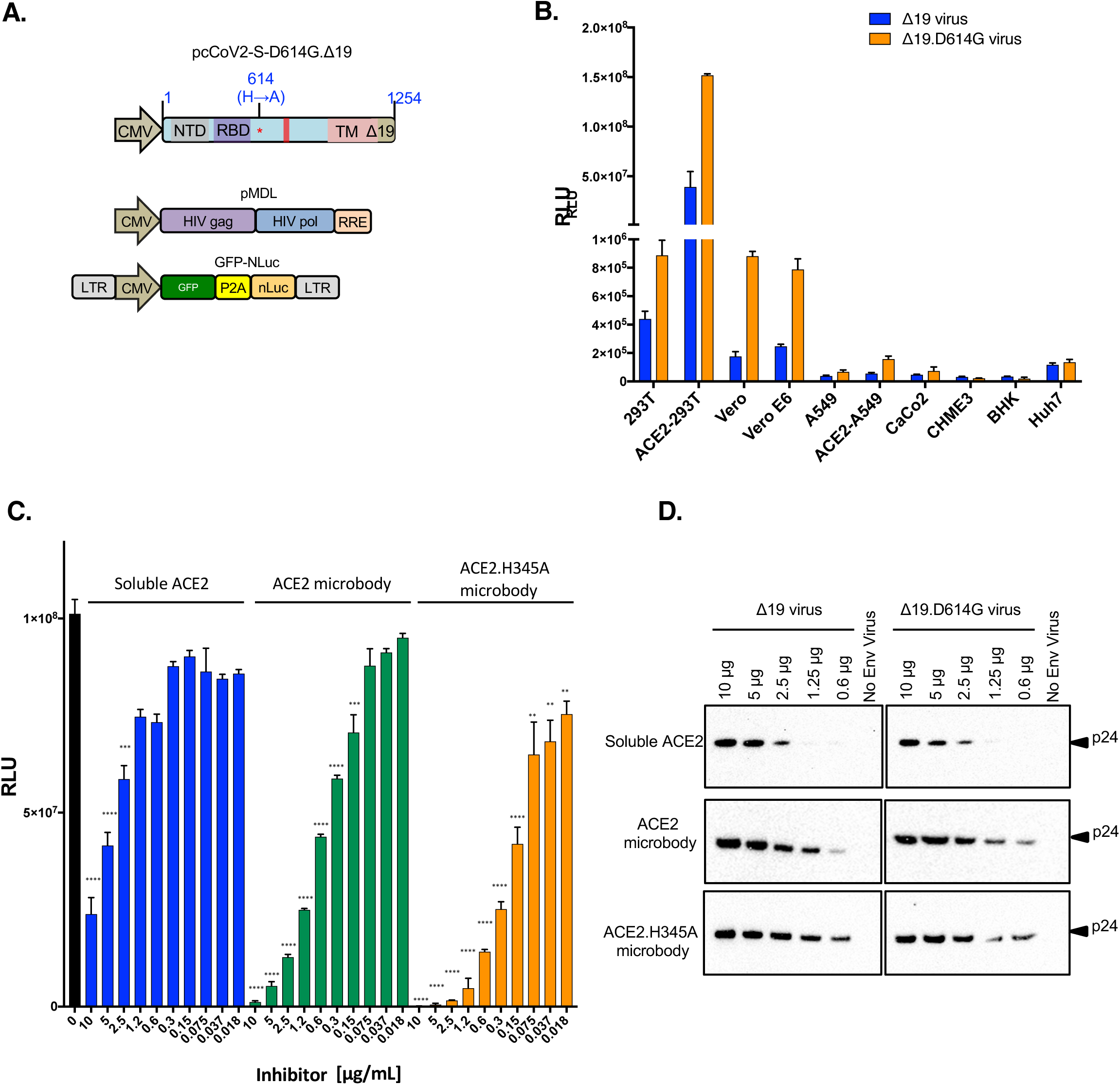
ACE2 microbody blocks entry of the D614G variant S protein pseudotyped virus infection. (A) The domain structure of the SARS-CoV-2 D614G Δ19 S expression vector is diagrammed above. Red star indicates the D614G mutation in the S protein. (B) A panel of cell-lines was infected with equivalent amounts of wild-type and D614G Δ19 S protein pseudotyped virus. (C) Serially diluted soluble ACE2 and ACE2 microbody proteins were mixed with D614G Δ19 S protein pseudotyped virus and added to target cells. Luciferase activity was measured 2 days post-infection. The data are shown as the mean of triplicates ± SD. The statistical significance of the data was calculated with the student-t test. (D) Ni-NTA agarose beads were coated with serially diluted soluble ACE2 and ACE2 microbody proteins. Wild-type and Δ19 S protein pseudotyped virions were added and allowed to bind. Unbound virions were removed after 30 min and the bound virions were detected by immunoblot analysis with anti-p24 antibody.

### ACE2 microbody is effective against other β coronavirus S proteins

To determine how well the ACE2 microbody would block the entry of other β coronaviruses, lentiviral virions pseudotyped by S proteins from a panel of different lineage 2 β coronaviruses that use ACE2 for entry (Letko and Munster, 2020) (pcSARS-CoV, pcSARS-CoV2, pcWIV1, pcLYRa11, pcRs4231, pcRs4084 and pcSHC014) were generated. The pseudotypes were incubated with soluble ACE2 and the ACE2 microbody proteins and their infectivity was then measured on ACE2.293T cells. The analysis showed that the ACE2 and H345A microbody proteins blocked all of the β coronavirus pseudotyped viruses while the antiviral activity of soluble ACE2 was significantly diminished in comparison (**Figure 7**). The results demonstrate the broad activity of the ACE2 microbody.

**Figure 7.**
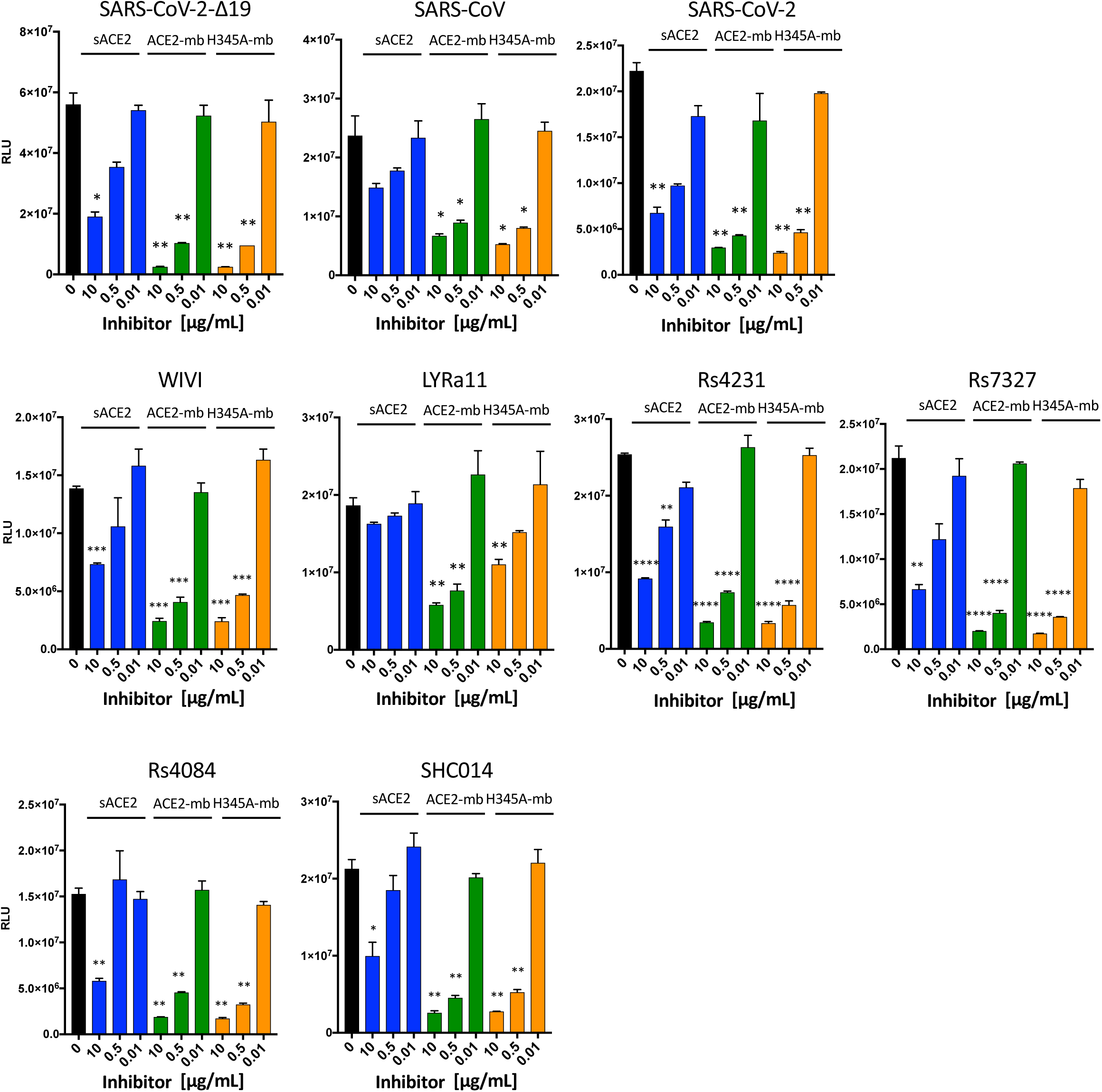
ACE2 microbody is effective against ACE2-using β coronavirus S proteins. Lentiviral virions pseudotyped by β coronavirus lineage 2 S proteins were treated with serially diluted soluble ACE2 proteins and then used to infect ACE2.293Tcells. The identity of the virus from which the S protein RBD is derived is indicated above each histogram. Luciferase activity was measured after 2 days. The data are displayed as the mean ± SD and significance is determined by student-t tests.

## Discussion

We report the development of a soluble form of ACE2 in which the ectodomain of ACE2 is fused to a single domain of the IgG heavy chain Fc. The domain renders the protein smaller than those fused to the full-length Fc yet retains the cysteine residues required for dimerization and the ability to increase the *in vivo* half-life (Maute et al., 2015). The microbody protein was shown to be a disulfide-bonded dimer in contrast to soluble ACE2 lacking the Fc domain which was dimeric but not nondisulfide-bonded. Although both proteins are dimeric, the ACE2 microbody had about 10-fold more antiviral activity than soluble ACE2 and bound to virions with a >4-fold increased affinity.

While high affinity anti-spike RBD monoclonal antibodies that potently inhibit SARS-CoV-2 infection will be of great value in the treatment of COVID-19, the soluble receptor proteins have advantageous features. The ACE2 microbody is of fully human origin so should be relatively non-immunogenic. In addition, it is expected to be broadly active against mutated variant spike proteins that may arise in the human population. The microbody was fully active against virus with the D614G S protein, a variant of increasing prevalence with increased infectivity *in vitro* (**Figure 6**) and was highly active against ACE2-specific S proteins from other β coronaviruses. It has been previously shown that a recombinant ACE2-Fc fusion had a major effect on blood pressure in a mouse model (Liu et al., 2018). It was therefore important to inactivate ACE2 carboxypeptidase activity in microbody to decrease unwanted effects on blood pressure associated with its use therapeutically. The H345A mutation alters one of the histidine essential for ACE2 catalytic activity yet did not impair antiviral activity against SARS CoV-2 or other β coronavirus spike proteins. In some of our analyses, the ACE2.H345A microbody appeared to be more active than the wild-type protein although the significance of this difference was unclear as the two proteins had similar activity in the live virus replication assay.

Escape from inhibition studies provided insight into the kinetics of virus infection and the mechanism of inhibition by the soluble receptors. Pretreatment of virus with the ACE2 microbody potently neutralized the virus as did simultaneous treatment addition of virus and microbody to cells. Furthermore, the protein retained its ability to prevent infection even when added to the culture at times after addition of virus, blocking infection by about 50% when added 1 hour after virus addition. The ACE2 microbody was partially active even on virus that had already attached to the cell. When virus was pre-bound for 2 hours, a time at which about 10% of the infectious virus had bound the cell, the ACE2 microbody retained the ability to prevent infection of about 50% of the bound virus (**Figure 5A**). Taken together, the experiments suggest a series of events in which the virus binds to cells over a period of about 4 hours. During this time, the ACE2 microbody is highly efficient, neutralizing nearly all of the free virus. Once the virus binds to the cell, the ACE2 microbody retains its ability to block infection for about 30 min, suggesting that binding is initially mediated by a small number of S proteins and that over 2 hours, additional S proteins are recruited to interact with target cell ACE2, a period during which the ACE2 microbody remains able to block the viral fusion reaction. Once a sufficient number of S protein:ACE2 interactions have formed, the virus escapes neutralization.

It was surprising that the ACE2 microbody had more antiviral activity than soluble ACE2 as both proteins are dimeric. In addition, the ACE2 microbody protein showed somewhat better binding to virions than soluble ACE2. The reasons for these differences are not clear. It is possible that the disulfide bonds of the ACE2 microbody stabilize the dimer or that they position the individual monomers in a more favorable conformation to bind to the individual subunits of the S protein trimer. It is worth noting that in most of the experiments, we used ACE2.293T cells that overexpress ACE2 compared to the cell-lines tested. On untransfected 293T cells that express barely detectable levels of ACE2, the antiviral activity of the microbody protein was increased, suggesting that the antiviral activity of the ACE2 microbody may be under-estimated by the use of ACE2 over-expressing cells.

Recent reports have described similar soluble ACE2 proteins. Recently soluble ACE2-related inhibitor including rhACE2 was shown to partially block infection (Case et al., 2020; Lei et al., 2020; Monteil et al., 2020) although the proteins had a short half-life (Wysocki et al., 2010) (< 2 hours in mice), limiting their clinical usefulness. In contrast a dimeric rhesus ACE2-Fc fusion protein had a half-life in mice in plasma greater than 1 week (Liu et al., 2018). The half-life of the ACE2 microbody *in vivo* has not yet been tested, but the protein retained antiviral activity for several days in tissue culture, significantly longer than longer than soluble ACE2 (**Figure S3**).

The phenomenon of antibody-dependent enhancement is caused by the interaction of the Fc domain of non-neutralizing antibody with the Fc receptor on cells which then serves to promote rather than inhibit virus neutralization. A similar phenomenon is possible with receptor-Fc fusion proteins by interaction with Fc receptor on cells. Because the ACE2 microbody contained only a single Fc domain, it was not expected to interact with Fc receptor. To test whether this was the case, we tested the ACE2 microbody in an enhancement assay using U937 cells which express Fc receptors. The ACE2 microbody protein did not detectably bind to cells that express the Fc γ receptor and the cells did not become infected, suggesting that this mechanism is not likely to play a role *in vivo* (**Figure S4 and not shown).**

Pseudotyped viruses have been highly useful for studies of SARS-CoV-2 entry. Vectors for producing SARS-CoV-2 lentiviral pseudotypes have been developed by several laboratories (Crawford et al., 2020; Nie et al., 2020; Ou et al., 2020; Schmidt et al., 2020; Shang et al., 2020; Xia et al., 2020). The vectors we report here produce pseudotyped lentiviral viruses with very high infectivity. The high infectivity of the pseudotypes produced is due in part to efficient expression of a codon-optimized Δ19 S protein and the efficient virion incorporation that results from the cytoplasmic tail truncation. The Δ19 S protein was present at only slightly higher levels on the cell surface than the full-length protein, suggesting that this small increase does not fully account for the large increase in virion incorporation. A possible explanation is that the full-length cytoplasmic tail sterically hinders virion incorporation by conflicting with the underlying viral matrix protein and that the deletion removes the conflict. Also, contributing to high viral titers, is the use of separate Gag/Pol packaging vector and lentiviral transfer vector as opposed to a lentiviral proviral DNA encoding Gag/Pol and the reporter gene, a strategy that resulted in higher reporter gene expression as shown in a direct comparison (not shown). Moreover, the dual luciferase/GFP reporter allows for measurement of infectious virus titer by flow cytometry and the high sensitivity of nanoluciferase read-out. The lentiviral pseudotypes are highly useful for rapidly titering neutralizing antibody in patient serum. In a study of over 100 sera from recovering patients, we found that the pseudotype assay results to be highly correlated with those of a live virus neutralization assay (submitted).

A feature of soluble receptors is that because the virus spike protein needs to conserve receptor binding affinity to maintain transmissibility, they should maintain their ability to neutralize S protein variants. SARS-CoV-2 S variants have been found to be circulating in the human population and it is likely that others are yet to emerge, some of which may be less sensitive to neutralization by the therapeutic monoclonal antibodies currently under development. The recently identified SARS-CoV-2 variant encoding the D614G S protein has been found to be spreading with increased frequency in the human population (Daniloski et al., 2020; Eaaswarkhanth et al., 2020; Korber et al., 2020; Zhang et al., 2020). The D614G S protein was found to be more resistant to shedding from the virion and to adopt a conformation that favors ACE2-binding and is in a more fusion-competent state (Yurkovetskiy et al., 2020; Zhang et al., 2020). We confirmed the increased infectivity of virions and find that the D614G S protein has a higher affinity for ACE2 as measured in a virion binding assay. Nevertheless, the ACE2 microbody maintained its ability to neutralize D614G S protein pseudotyped virus. The ability of the ACE2 microbody to neutralize diverse β coronaviruses suggest that it may also be able to neutralize novel ACE2 using coronaviruses that may be transferred to the human population in the future. The microbody protein could serve as an off-the-shelf reagent that could be rapidly deployed.

## Author Contributions

N.R.L., T.T. and C.N. designed the experiments. T.T. and NRL. wrote the manuscript. T.T., C.F., R.K. and K.S. did the experiments. H.B.G. did the SEC/MALs and P.B. provided expertise and guidance.

## Acknowledgements

We thank Lin Xinhua and Hanna Nazeeh (NYULH) and Benjamin tenOever for cell-lines and Michael Letko and Vincent Munster (NIH) for the β coronavirus spike protein expression vectors. The work was funded by grants from the NIH to N.R.L. (DA046100, AI122390 and AI120898) and to P.B.J in support of H.B.G. (P01-AI38398-S1).

## Supplemental Information

**Figure S1.**
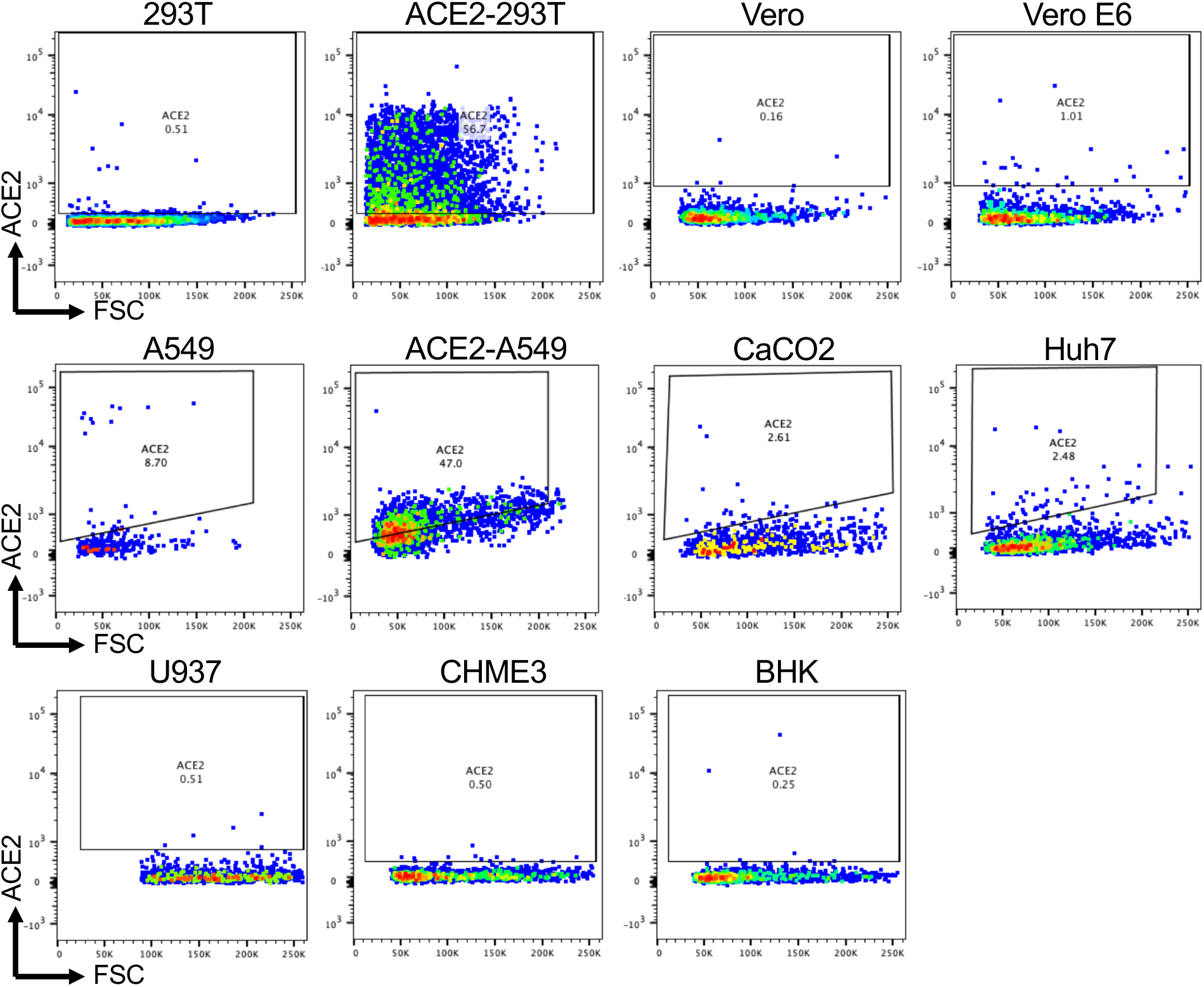
ACE2 expression levels in cell-lines. (A) The indicated cell lines were stained with anti-ACE2 antibody and Alexa fluor 594-conjugated anti-mouse IgG secondary antibody and analyzed by flow cytometry.

**Figure S2.**
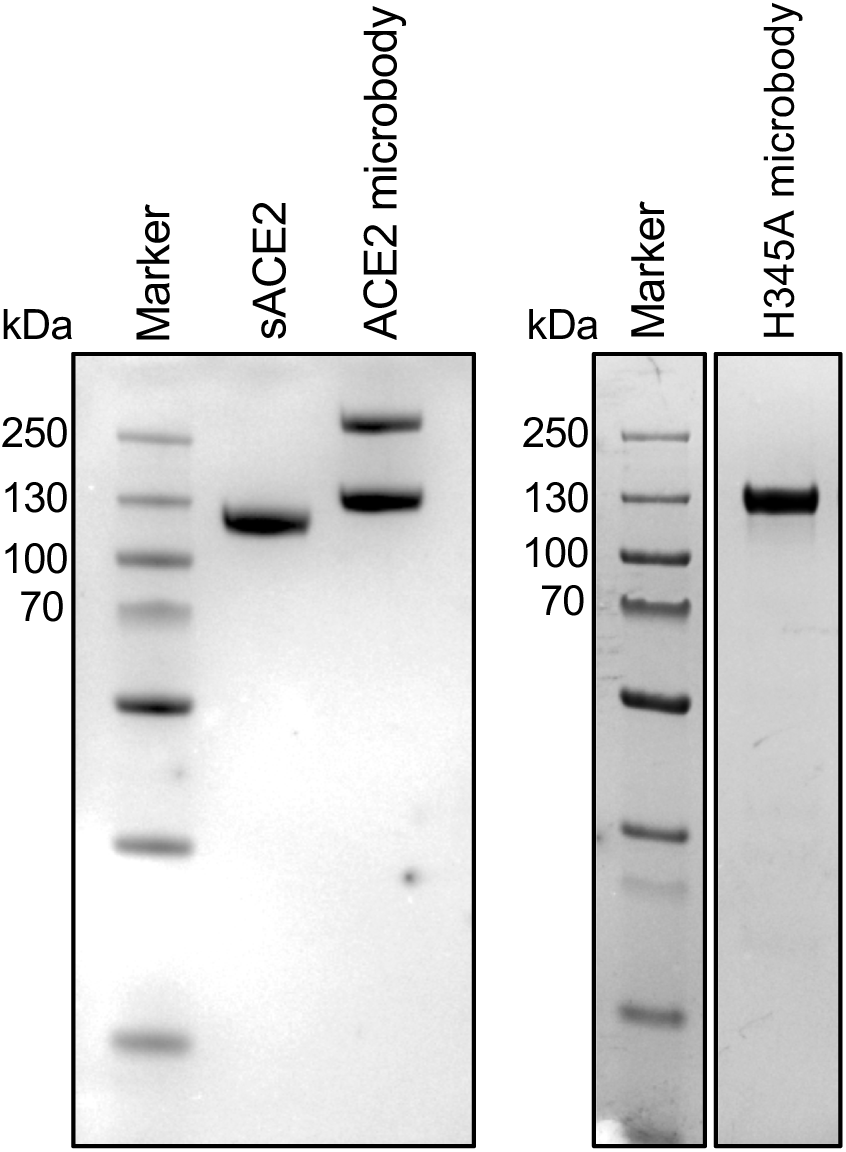
SDS-PAGE analysis of purified proteins. 30 μg of soluble ACE2, ACE2 microbody (left) and ACE2.H345A microbody (right) were analyzed by Coomassie blue stained SDS-PAGE under reducing conditions. Note, the ACE2 microbody dimer is partially resistant to reduction.

**Figure S3.**
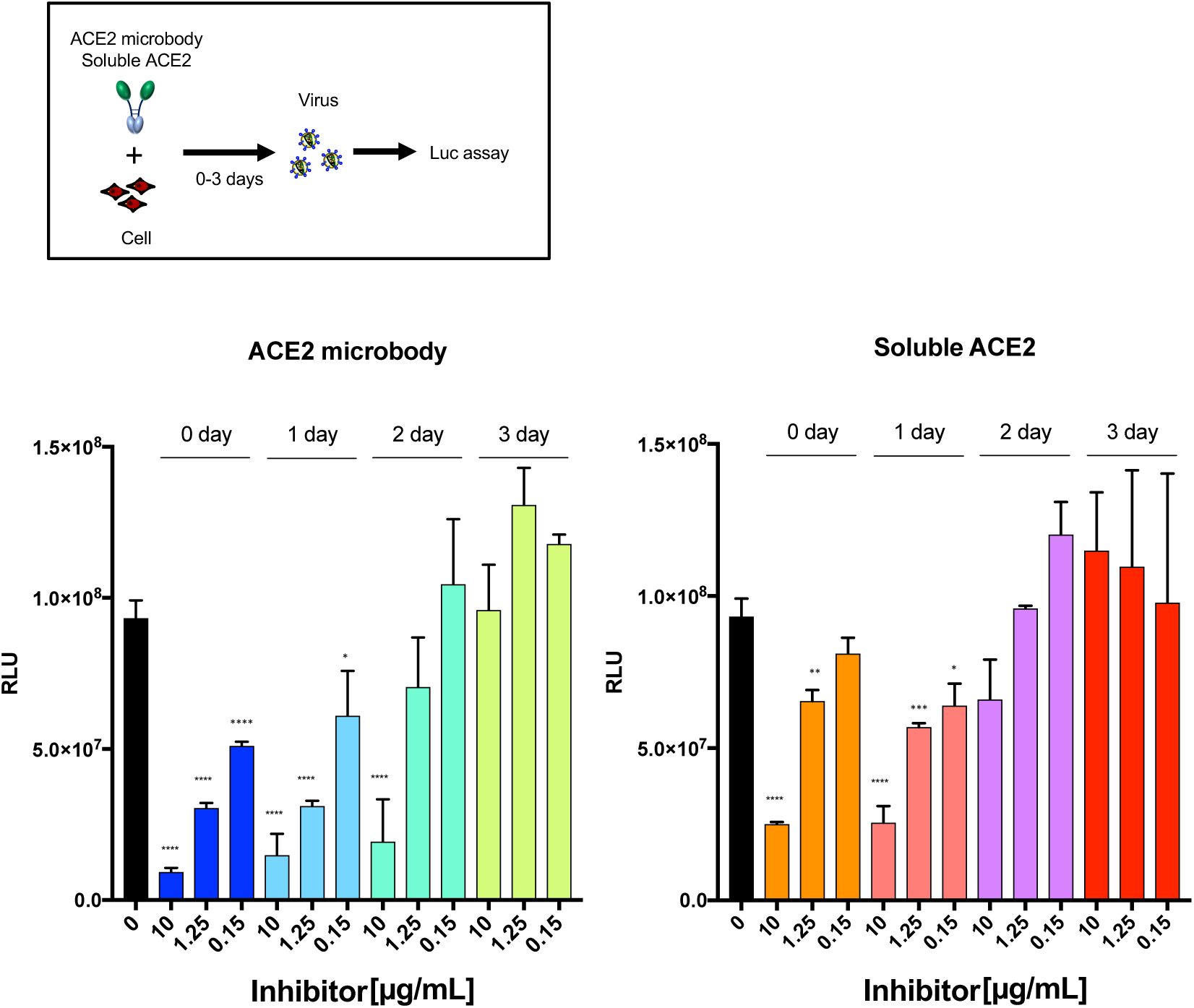
Increased stability of the ACE2 microbody. Serially diluted sACE2 and ACE2 microbody proteins were added to ACE2.293T target cell cultures. The cells were infected either immediately with SARS-CoV-2 pseudotyped lentivirus or 1, 2 or 3 days later. Luciferase activity was measured 2 days post-infection. The data are presented as the mean of triplicates ± SD. Statistical significance was calculated by the student-t test.

**Figure S4.**
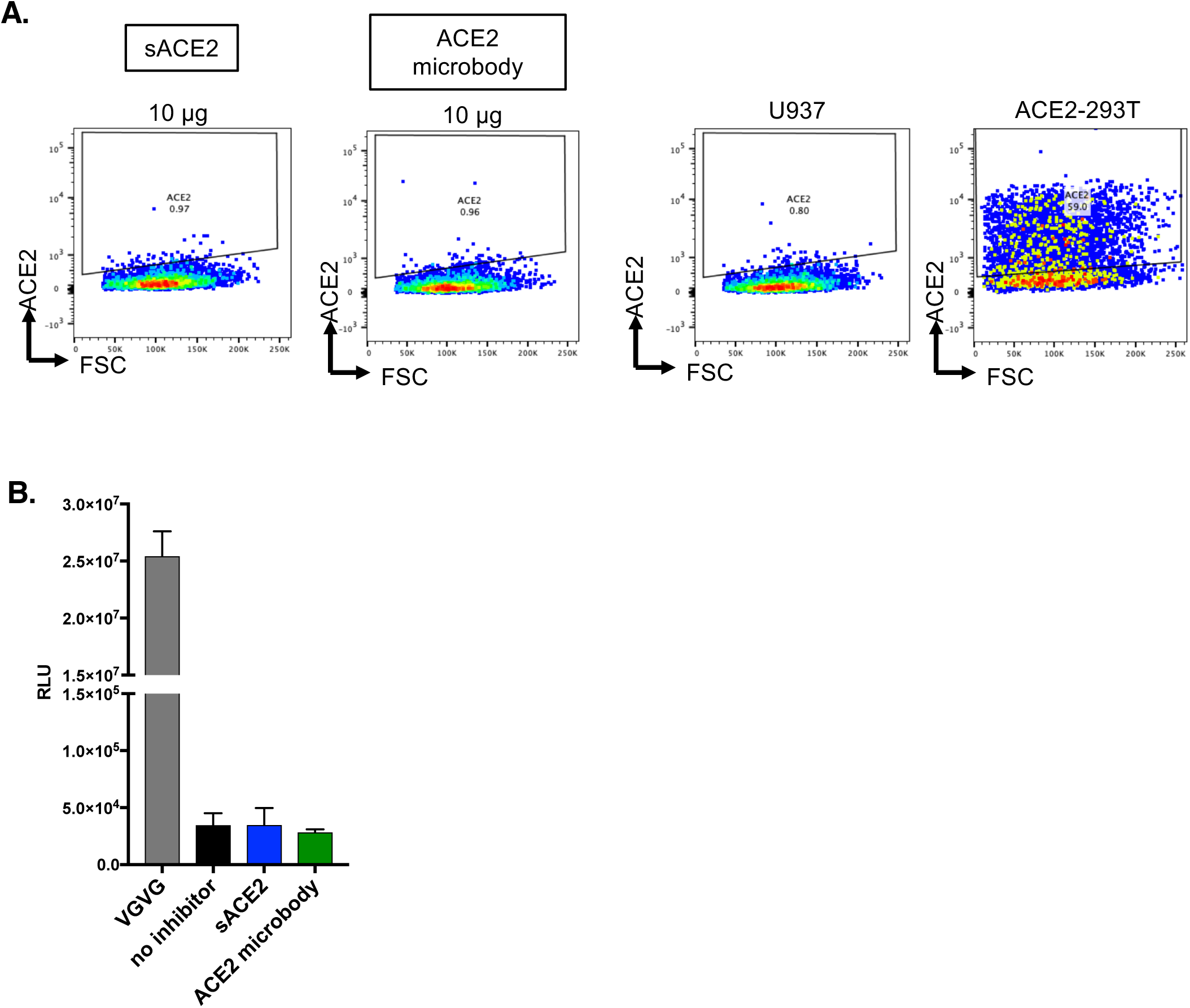
ACE2 microbody does not bind to Fc receptors. (A) U937 cells were incubated for 30 min. with serially diluted soluble ACE2 or ACE2 microbody. Unbound soluble ACE2 proteins were removed and cell surface-bound proteins were detected by flow cytometry with anti-ACE2 antibody and Alexa fluor 594-conjugated anti-mouse IgG secondary antibody. ACE2.293T cells were analyzed as a positive control for ACE2 staining.

**Figure S5.**
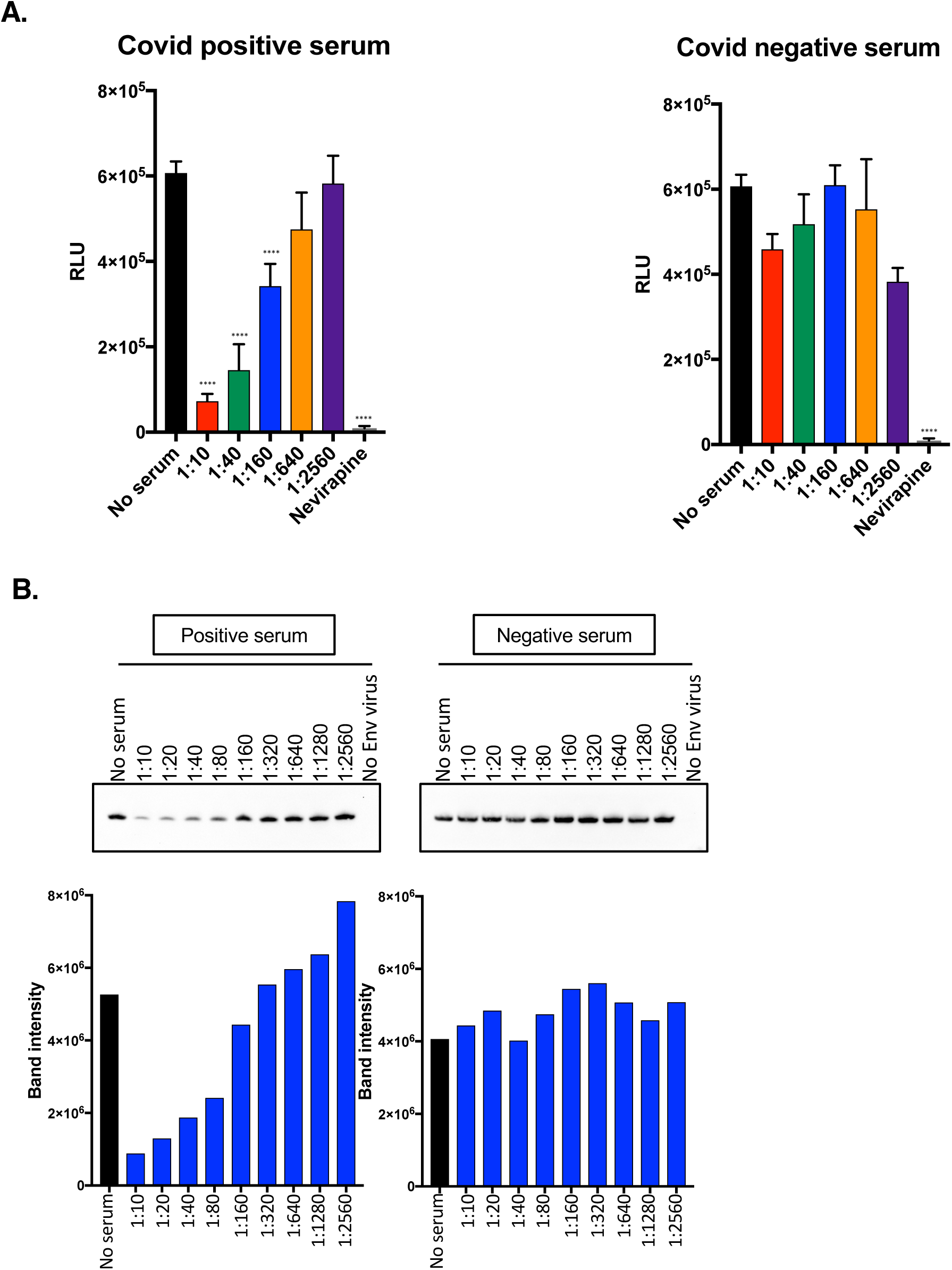
Titer of convalescent patient serum with SARS-CoV-2 lentiviral pseudotyped virus and effect of serum in virion binding assay. (A) Serially diluted serum from a COVID-19 patient (left) and healthy donor (right) was incubated for 30 minutes with pseudotyped virus and then added to Vero E6 cells. Two days post-infection, luciferase activity was measured. Nevirapine was added to one sample to confirm that signals were the result of bone fide infection. The data are displayed as the mean of triplicates ± SD. Statistical significance was calculated by the student-t tests. (B) The ability of convalescent patient serum to block virus binding to ACE2 was tested. Ni-NTA agarose beads were coated with ACE2 microbody proteins. Serially diluted convalescent patient serum or healthy donor serum was incubated for 30 minutes with pseudotyped virions. The virions were then incubated for 1 hour with ACE2 microbody coated-beads. Free virions were removed and the bound protein was analyzed on an immunoblot probed with anti-p24 antibody. A histogram showing band intensities is shown below.

## Methods

### Plasmids

The dual GFP/nanoluciferase lentiviral vector pLenti.GFP.NLuc was generated by overlap extension PCR. A DNA fragment encoding GFP was amplified with a forward primer containing a BamH-I site and a reverse primer encoding the P2A sequence. The nanoluciferase gene (NLuc) was amplified with a forward primer encoding the P2A motif and a reverse primer containing a 3’-Sal-I site. The amplicons were mixed and amplified with the external primers. The fused amplicon was cleaved with BamH-I and Sal-I and cloned into pLenti.CMV.GFP.puro (Addgene plasmid #17448, provided by Eric Campeau and Paul Kaufman) (Campeau et al., 2009).

The SARS-CoV-2 S expression vector pcCOV2.S was chemically synthesized as DNA fragments A and B encoding codon-optimized 5’ and 3’ halves, respectively, of the S gene of Wuhan-Hu-1/2019 SARS-CoV-2 isolate (Table S2 and S3). Fragment A was amplified with a forward primer containing a Kpn-I site and reverse primer containing an EcoR-I site. The amplicon was cleaved with Kpn-I and EcoR-I and cloned into pcDNA6 (Invitrogen). Fragment B was amplified with a forward primer containing an EcoR-I site and reverse primer containing Mlu-I and Xho-I sites. The amplicon was cleaved with EcoR-I and Xho-I and cloned into pcDNA6. The cloned fragment A was then cleaved with Mlu-I and Xho-I and cloned into the Mlu-I and Xho-I sites in the fragment B-containing plasmid. To generate the SARS-CoV-2 S Δ19 expression vector pcCoV2.S.Δ19, the codon-optimized S gene was amplified with a forward primer containing a Kpn-I site and reverse primer that deleted the 19 carboxy-terminal amino acids and contained an Xho-I site. The amplicon was cloned into the Kpn-I and Xho-I of pcDNA6. The D614G mutation in S was generated by overlap extension PCR of the Δ19 S gene using internal primers overlapping the sequence encoding D614G and cloned into pcDNA6. Beta coronavirus spike expression vectors (Letko and Munster, 2020) (pcSARS-CoV, pcSARS-CoV2, pcWIV1, pcLYRa11, pcRs4231, pcRs4084 and pcSHC014) were kindly provided by Michael Letko and Vincent Munster (NIH).

ACE2 expressing lentiviral vector pLenti.ACE2 was generated by amplifying an ACE2 cDNA (Origene) with a forward primer containing an Xba-I site and reverse primer containing a Sal-I site. The amplicon was cleaved with Xba-I and Sal-I and cloned into pLenti.CMV.GFP.puro in place of GFP. The soluble ACE2 expression vector pcsACE2 was generated by amplifying the extracellular domain of ACE2 with a forward primer containing a Kpn-I site and reverse primer encoding an in-frame 8XHis-tag and Xho-I site. The amplicon was then cloned into the Kpn-I and Xho-I sites of pcDNA6. The ACE2 microbody expression vector pcACE2-microbody was generated by overlap extension PCR that fused the extracellular domain of ACE2 with human immunoglobulin G heavy chain Fc domain 3 using a forward primer containing a Kpn-I site and reverse primer containing an 8XHis-tag and Xho-1 site. The amplicon was cloned into the Kpn-I and Xho-I sites of pcDNA6. Expression vector pcACE2.H345A-microbody that expressed the ACE2.H345A microbody was generated by overlap extension PCR using primers that overlapped the mutation. Full-length cDNA sequence, primer sequences and amino acid sequences are shown in Tables S1-3.

### Human sera

Control and recovered patient sera were collected from patients through the NYU Vaccine Center with written consent under I.R.B. approval (IRB 20-00595 and IRB 18-02037) and were deidentified.

### Cells

Vero E6, CaCO2, A549, ACE2 A549, BHK, Huh7 293T, Vero and CHME3 cells were cultured in Dulbecco’s modified Eagle medium (DMEM) supplemented with 10% fetal bovine serum (FBS) and penicillin/streptomycin (P/S) at 37°C in 5% CO2. CaCO2 cells were cultured in DMEM/10% FBS/1% nonessential amino acids. U937 cells were cultured in RPMI/10% FBS/ with P/S. ExpiCHO-S (Thermo Fisher Scientific) were cultured in ExpiCHO expression medium at 37 °C in 8% CO2. Cell-line ACE2 expression levels were quantified by staining with anti-ACE2 antibody (NOVUS) and Alexa-fluor 594-conjugated goat anti-mouse IgG (Biolegend) and pacific blue viability dye. Data were analyzed by flow cytometry with Flowjo software. ACE2.293T cells were established by lipofection of 293T cells with pLenti.ACE2-HA using lipofectamine 2000 (Invitrogen). After 2 days, the cells were selected in 1 μg/ml puromycin and cloned at limiting dilution. Single cell clones were expanded and analyzed by flow cytometry and a single clone was chosen.

### SARS-CoV-2 pseudotype reporter virus assay

SARS-CoV-2 S protein pseudotyped lentiviral stocks were produced by cotransfecting 293T cells (4 × 10^6^) with pMDL, pLenti.GFP-NLuc, S protein expression vector and pRSV.Rev at a mass ratio of 4:3:4:1 by calcium phosphate coprecipitation. S protein expression vectors used were pcCoV2.S, pcCoV2.S-Δ19 or the β coronavirus RBD expression vectors. Control viruses were produced substituting the S protein vector for pcVSV or with pcDNA6 to produce virus lacking S protein. Virus-containing supernatant was harvested 2 days post-transfection, passed through a 0.45 μm filter and concentrated by ultracentrifugation over a 20% sucrose cushion at 30,000 RPM for 90 min in an SW40.1 rotor in a Beckman Optima L-100K ultracentrifuge (Brea, CA). The pellet was resuspended to 1/10 the initial volume in DMEM/10% FBS and frozen in aliquots at −80°C. Virus stocks were titered on 293T by flow cytometry and for luciferase activity. The p24 concentration was measured and the virus was used at a concentration of 1.0 μg/ml. To test the inhibitory activity of soluble receptors and convalescent sera, 50 μl serially diluted inhibitor or convalescent patient serum was incubated for 30 min at room temperature with 5 μl pseudotyped reporter virus (approximately 5 × 10^6^ cps luciferase activity/μl) at a MOI of 0.1 in a volume of 100 μl. The mixture was added to ACE2.293T cells in a 96 well tissue culture dish containing 1 × 10^4^ cells/well. After 2 days, the culture medium was removed and 50 μls Nano-Glo luciferase substrate (Promega) and 50 μls medium was added to each well. The supernatant (70 μls) was transferred to a microtiter plate and the luminescence was read in an Envision 2103 microplate luminometer (PerkinElmer). Alternatively, the GFP+ cells were quantified by flow cytometry with pacific blue viability dye to exclude dead cells (Biolegend).

### Protein purification and molecular mass determination

293F cells (Thermo Fisher) at a density of 2.5 × 10^6^ cells/ml were transfected with microbody expression vector plasmid DNA using polyethyleneimine (Polysciences, Inc) at a 1:3 plasmid:PEI ratio. The cells were then cultured at 30°C and at 12 hours post-transfection 10 mM sodium butyrate was added. After 4 days, the supernatant culture medium was collected, filtered and adjusted pH to 8.0. The medium was passed over a 5 ml HiTrap Chelating column charged with nickel (GE healthcare), washed with 30 ml of buffer containing 20 mM Tris pH 8, 150 mM NaCl, 10 mM imidazole and the bound protein was eluted in buffer containing 250 mM imidazole. The eluate was concentrated to 1.0 ml and loaded onto a Superdex 200 size-exclusion column (GE healthcare) in running buffer containing 10 mM Tris pH 7.4, 150 mM NaCl. Protein containing fractions were pooled and concentrated. The purified proteins were analyzed on a 4-12% Bis-Tris SDS-PAGE stained with Coomassie blue.

The absolute molecular masses of the purified protein complexes were determined by SEC/MALs. The proteins were injected onto a Superdex 200 10/300 GL gel-filtration chromatography column equilibrated in sample buffer that was connected to a Dawn Heleos II 18-angle light-scattering detector (Wyatt Technology), a dynamic light-scattering detector (DynaPro Nanostar; Wyatt Technology) and an Optilab t-rEX refractive index detector (Wyatt Technology). The data were collected at 25°C at a flow rate of 0.5 ml/minute every second. The molecular mass of each protein was determined by analysis with ASTRA 6 software.

### Virion pull-down assay

293T cells were transfected by lipofection with 4 μg pcACE2-microbody. At 72 hours post-transduction, 0.5 ml of culture supernatant was incubated with nickel-nitrilotriacetic acid-agarose beads (QIAGEN). The beads were washed, and bound protein was eluted with Laemmle loading buffer. The proteins were analyzed on an immunoblot probed with mouse anti-6XHis antibody (Invitrogen) and horseradish peroxidase (HRP)-conjugated goat anti-mouse IgG secondary antibody (Sigma-Aldrich). The proteins were visualized using luminescent substrate and scanned on a LI-COR Biosciences FC Imaging System (LI-COR Biotechnology). Ratios were calculated as the His (spike) signal intensity divided by the p24 signal intensity for an identical exposure of the blot.

### Immunoblot analysis

Transfected cells were lysed in buffer containing 50 mM HEPES, 150 mM KCl, 2 mM EDTA, 0.5% NP-40, and protease inhibitor cocktail. Protein concentration in the lysates was measured by bicinchoninic protein assay and the lysates (40 μg) were separated by SDS-PAGE. The proteins were transferred to polyvinylidene difluoride membranes and probed with anti-HA mAb (Covance), mouse anti-His mAb (Invitrogen) and anti-GAPDH mAb (Life Technologies) followed by goat anti-mouse HRP-conjugated second antibody (Sigma). The blots were visualized using luminescent substrate (Millipore) on a LI-COR Bio-sciences FC Imaging System.

### Binding assay

Soluble ACE2 proteins (10 μg) were mixed with 20 μl nickel beads for 1 hour at 4°C. Unbound protein was removed by washing the beads with PBS. The beads were resuspended in PBS and mixed with 40 μl pseudotyped lentiviral virions After 1 h incubation at 4°C, the beads were washed with PBS and resuspended in reducing Laemmli loading buffer and heated to 90°C. The eluted proteins were separated by SDS-PAGE and analyzed on an immunoblot probed with anti-p24 antibody (AG3.0) followed by goat anti-mouse HRP-conjugated second antibody.

### Live SARS-CoV-2 neutralization assay

mNeonGreen SARS-CoV-2 (Xie et al., Cell Host and Microbe 2020) was obtained from the World Reference Center for Emerging Viruses and Arboviruses at the University of Texas Medical Branch. The virus was passaged once on Vero E6 cells (ATCC CRL-1586), clarified by low-speed centrifugation, aliquoted, and stored at −80°C. The infectious virus titer was determined by plaque assay on Vero E6 cells after staining with crystal violet. Virus neutralization was determined as previously described (Xie et al, bioRxiv 2020). ACE2.293T cells were seeded in a 96-well plate (1 × 10^4^/well). The next day, mNeonGreen SARS-CoV-2 (MOI = 0.5) was mixed 1:1 with serially 2-fold diluted soluble ACE2 protein in DMEM/2% FBS and incubated for 1 hour at 37°C. The virus:protein mixture was then added to the ACE2 cells and incubated for 24 hours. at 37°C in 5% CO2. The cells were fixed with 4% paraformaldehyde, stained with DAPI and the mNeonGreen+ cells were counted on a CellInsight CX5 Platform high content microscope (Thermo Fisher).

### Data analysis and statistics

All experiments were performed in technical duplicates or triplicates and data were analyzed using GraphPad Prism (Version 7 7.0e). Statistical significance was determined by the two-tailed, unpaired t test. Significance was based on two-sided testing and attributed to p< 0.05. Confidence intervals are shown as the mean ± SD or SEM. (*P≤ 0.05, **P≤ 0.01, ***P≤0.001, ****P≤0.0001).

---

**Table S1.**
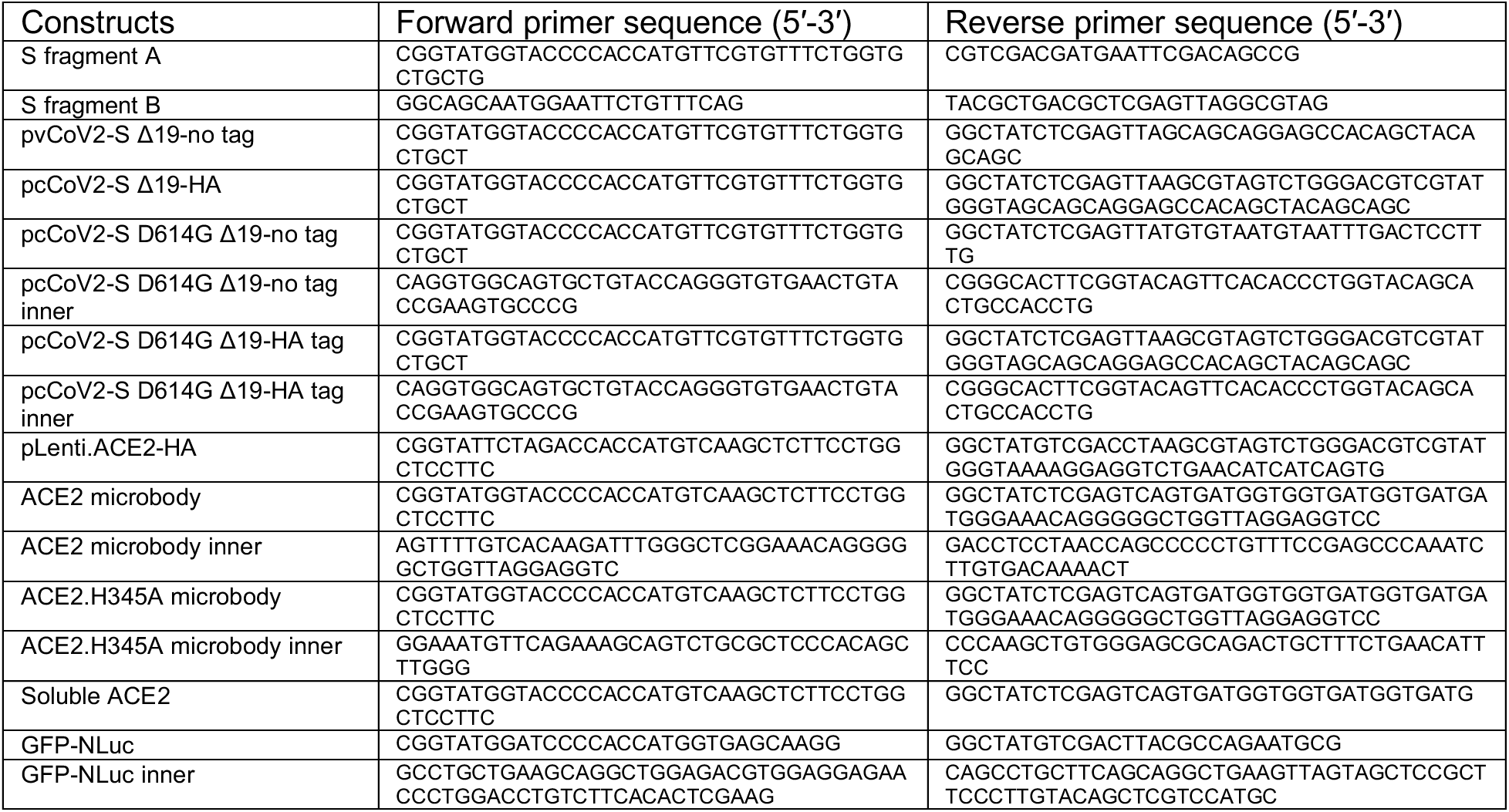
Primer sequence

**Table S2.**
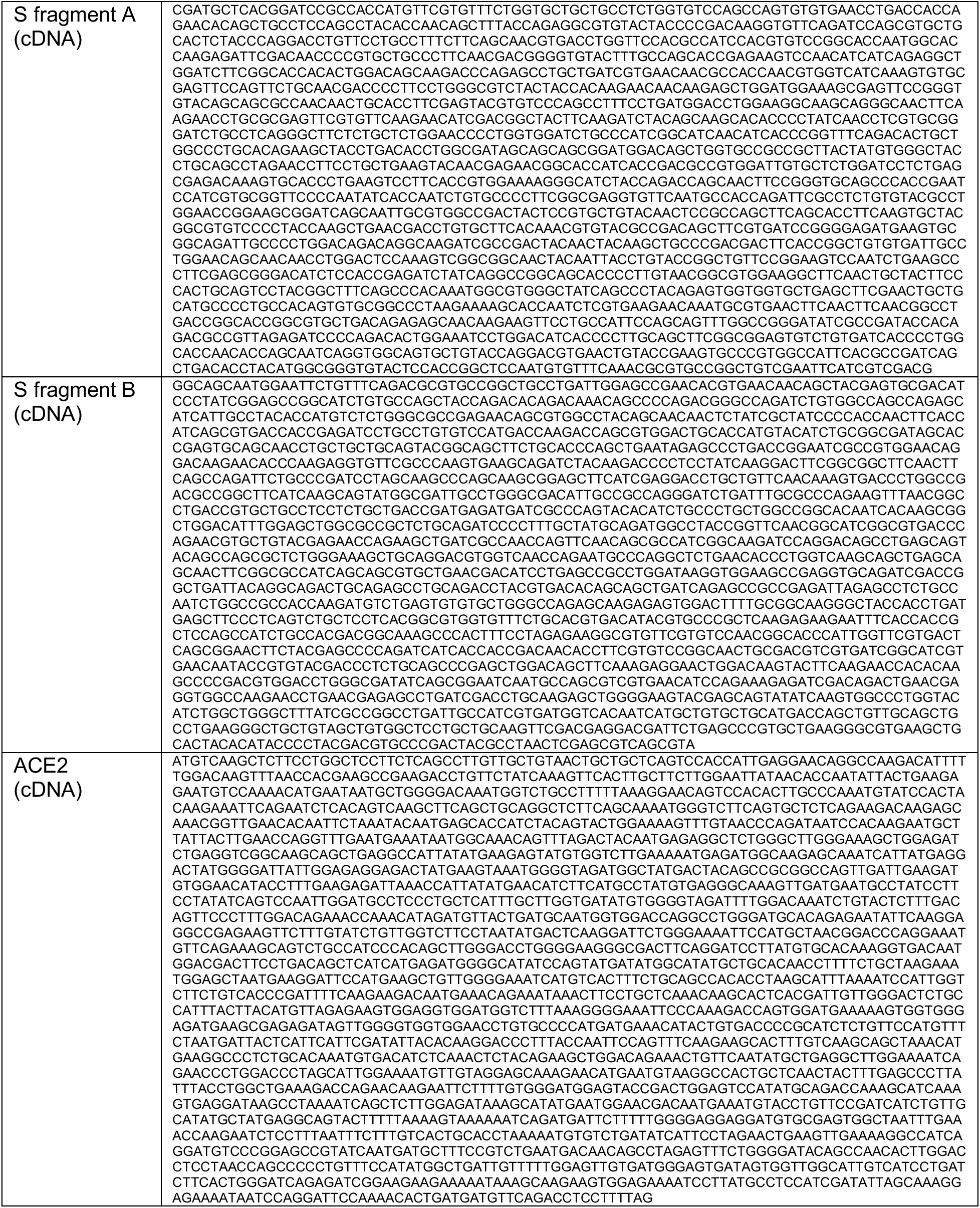
cDNA sequence

**Table S3.**
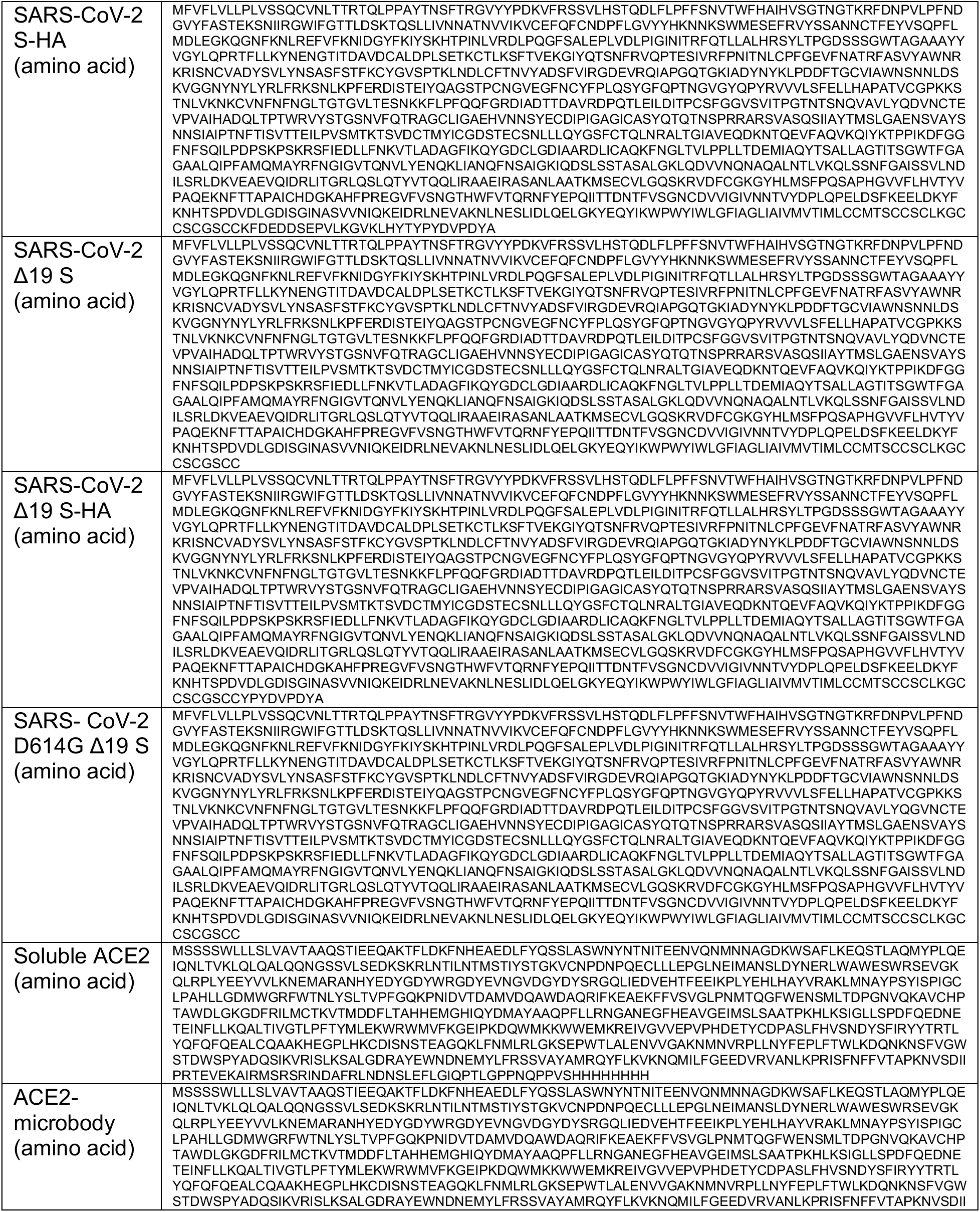
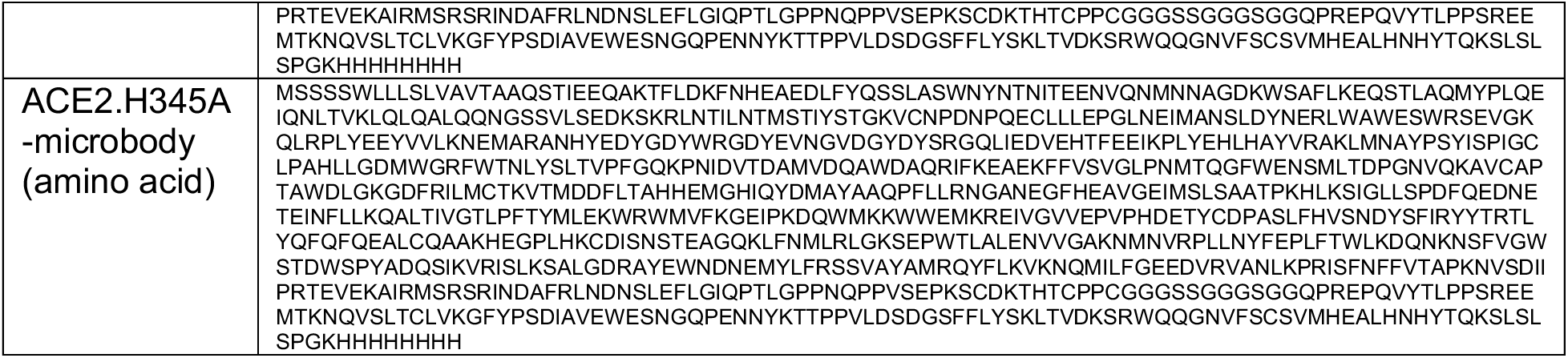
Amino acid sequence

